# CDK8 fine-tunes IL-6 transcriptional activities by limiting STAT3 resident time at the gene loci

**DOI:** 10.1101/2020.03.19.998351

**Authors:** J. Martinez-Fabregas, L. Wang, E. Pohler, A. Cozzani, M. Kazemian, S. Mitra, I. Moraga

## Abstract

Cytokines are highly pleiotropic ligands that critically contribute to a balanced immune response. We have an incomplete understanding of how cytokines elicit their functional pleiotropy, which has limited their therapeutic use. Here, using Interleukin-6 (IL-6) as a model system, we have performed detailed phosphoproteomic and transcriptomic studies in human Th-1 cells to address the molecular bases defining cytokine functional pleiotropy. We have identified CDK8 as a new negative regulator of STAT3 transcriptional activities that contributes to the diversification of IL-6 responses. We found that CDK8 is a major regulator of the IL-6 phosphoproteome and interacts with STAT3 in the nucleus upon IL-6 stimulation. Inhibition of CDK8 activity, using specific small molecules inhibitors, reduced the IL-6-induced phosphoproteome by 23% in Th-1 cells, including STAT3 Ser727 phosphorylation. This, in turn, resulted in retention of tyrosine phosphorylated STAT3 in the nucleus, which increased the binding of STAT3 to target DNA sites in the genome with a concomitant increase in STAT3 mediated transcriptional activity. Importantly, inhibition of CDK8 activity under Th-17 polarizing conditions resulted in an enhancement of Th-17 differentiation. Our results support a model where CDK8 regulates STAT3 transcriptional processivity via modulation of its gene loci resident time, critically contributing to tuning STAT3 mediated responses.

## INTRODUCTION

Cytokines play a crucial role overseeing the correct functioning of the immune system (Lin and Leonard, 2019). In response to environmental changes immune cells secrete cytokines, which in turn determine the nature, potency and duration of the immune response (Arai et al., 1990). Despite the functional relevancy of this family of ligands, the molecular bases governing their large functional pleiotropy remains poorly defined. Cytokines exert their activities by dimerizing/oligomerizing surface receptors and triggering the tyrosine phosphorylation of STATs transcription factors via JAK kinases (Gorby et al., 2018; Martinez-Fabregas et al., 2019; Stroud and Wells, 2004; Wang et al., 2009; Wilmes et al., 2020). This in turn leads to the nuclear translocation of STATs and the induction of specific gene expression programs and bioactivities (Poli and Camporeale, 2015; Schindler et al., 2007). In addition to STATs, cytokines activate additional signaling pathways including the PI3K/Akt pathway, the MAPK pathway and the src kinase pathway (Ali et al., 2015; Han et al., 2012; Leonard and Lin, 2000; Liu et al., 2013; Silva, 2004; Zhang and Liu, 2002), but how these pathways contribute to cytokine functional pleiotropy is poorly understood. A systematic study defining cytokine signaling signatures and their role in promoting cytokine functional pleiotropy would be highly informative.

STATs can be modified in conserved tyrosine or serine residues (Decker and Kovarik, 2000). While STAT tyrosine phosphorylation plays a critical role in mediating cytokine responses, the role of STAT serine phosphorylation in cytokine mediated activities is more controversial (Chung et al., 1997; Decker and Kovarik, 2000; Kim and Baumann, 1997; Wen et al., 1995). Whereas some studies report a positive effect of STAT serine phosphorylation in driving STAT transcriptional activities, others have reported the opposite effect (Bancerek et al., 2013; Decker and Kovarik, 2000; Levy and Darnell, 2002; Lim and Cao, 1999; Steen et al., 2016; Wen et al., 1995; Yokogami et al., 2000). How STAT serine phosphorylation modulates the immune response has been highly overlooked, with only a handful of studies available, mainly focused on STAT1 serine phosphorylation and its role modulating the innate response (Pilz et al., 2003; Rhee et al., 2003; Varinou et al., 2003). In macrophages, CDK8 appears to be the main kinase driving STAT1 serine phosphorylation in response to IFN gamma stimulation (Bancerek et al., 2013). Blockage of STAT1 serine phosphorylation, using STAT1 serine mutants, significantly altered the IFN gamma transcriptional response and its ability to clear *Listeria monocytogenes* infection (Varinou et al., 2003). However, the role that serine phosphorylation plays in regulating the activities of other STATs in different immune cells is less well known.

IL-6 represents a classical paradigm for cytokine functional pleiotropy. IL-6 acts as a central regulator of the immune response by triggering both pro-inflammatory and anti-inflammatory responses (Hunter and Jones, 2015; O’Shea and Murray, 2008; Rose-John, 2018; Scheller et al., 2011). IL-6 drives inflammatory processes by modulating the adaptive and innate immunity arms. On the one hand, IL-6 promotes the differentiation of Th-17 cells, while inhibiting the differentiation of T regulatory cells (Jones et al., 2010; Korn et al., 2008). On the other hand, IL-6 recruits myeloid cells to sites of inflammation (Fielding et al., 2008; Gabay, 2006). IL-6 also elicits anti-inflammatory responses including muscle and liver regeneration, regulation of metabolic processes and modulation of pain (Mauer et al., 2015; Poli and Camporeale, 2015; Scheller et al., 2011). Additionally, dysregulation of IL-6 or IL-6 mediated responses is often associated with inflammatory disorders, making this cytokine highly relevant for human health (Jones and Jenkins, 2018; Tanaka et al., 2014). IL-6 exerts its activities by triggering the activation of the JAK1/STAT1/STAT3 signaling pathway upon recruitment of IL-6Ra and gp130 receptor subunits (Heinrich et al., 1998; Martinez-Fabregas et al., 2019; Servais et al., 2019). Moreover, non-JAK/STAT pathways have been described to be activated by IL-6 in a cell context manner (Mauer et al., 2015). The molecular bases that allow IL-6 to elicit its vast array of immune-modulatory activities are not known.

In this study we set to characterize how the signaling initiated by IL-6 in human T cells leads to its functional pleiotropy. Using an antibody array targeting 28 relevant signaling molecules, we found that IL-6 preferentially engages the JAK/STAT1/STAT3 pathway in T cells. We could detect both STAT3 tyrosine and serine phosphorylation in response to IL-6 stimulation, with STAT3 serine phosphorylation exhibiting a delayed activation. To gain further insights into the IL-6 signalosome in T cells, we performed an unbiased phosphoproteomic study. This study revealed that the majority of phosphopeptides modified by IL-6 treatment belonged to nuclear proteins, highlighting a critical role of this compartment in defining IL-6 responses. Using a battery of small molecule inhibitors, we identified CDK8 and CDK9 as the kinases driving serine phosphorylation of STAT3 upon IL-6 stimulation in T cells. Interestingly, inhibition of these kinases produced a more sustained STAT3 tyrosine phosphorylation and a longer STAT3 nuclear retention upon IL-6 stimulation, suggesting that STAT3 serine phosphorylation regulates STAT3 nuclear dynamics. Using proximity ligation studies, we confirmed the increased interaction between STAT3 and CDK8 and CDK9 in the nucleus upon IL-6 stimulation. Inhibition of the activity of these two kinases resulted in a more robust interaction with STAT3, even in the absence of IL-6 stimulation. ChIP-Seq and RNA-Seq studies revealed a global increase in STAT3 chromatin binding upon inhibition of CDK8, resulting in the induction of a larger gene expression program by IL-6. In agreement with this enhanced gene expression program, IL-6 induced a more robust differentiation of Th-17 cells upon CDK8 inhibition *in vitro*. Overall, our studies identify a new STAT3 regulatory mechanism, whereby CDK8 and CDK9 modulates STAT3 processivity by controlling its chromatin binding dwell time and transcriptional activity through serine phosphorylation. These observations open new venues to manipulate IL-6 and STAT3-mediated responses by fine-tuning CDK8/9 activities.

## RESULTS

### Interleukin-6 signaling preferentially induces tyrosine/serine phosphorylation of STAT1/3 in human T cells

IL-6 critically contributes to modulate the T cell response. Yet, we have a poor understanding of the signaling networks engaged by IL-6 in T cells, and their role in shaping IL-6 immune activities. To gain insight into the IL-6 signalosome in T cells, we carried out detailed signaling studies in human resting or activated CD4^+^ and CD8^+^ T cells stimulated with IL-6. IL-6 receptor expression varies significantly among different T cell populations and environmental contexts (Sup. Figure 1 and (Betz and Muller, 1998; Jones et al., 2010; Ridgley et al., 2019), making the study of IL-6 signaling in T cells challenging. To minimize this variability, we have used Hyper-IL-6 (HyIL-6) for our signaling studies. HyIL-6 is a synthetic heterodimer comprised of IL-6Ra and IL-6 proteins connected by a gly/ser linker (Fischer et al., 1997). HyIL-6 triggers signaling in all cells expressing gp130, producing a more robust and homogenous signaling output (Rose-John, 2012). Dose-response (Figure 1A) and kinetic signaling studies (Figure 1B) showed that resting and activated CD4^+^ and CD8^+^ T cells respond to HyIL-6 treatment by Tyr phosphorylating STAT1 and STAT3 transcription factors. The STAT1/3 activation amplitudes elicited by HyIL-6 in these cells however differs significantly, with activated CD4^+^ and CD8^+^ T cells triggering between 2-3 folds higher STAT1/3 phosphorylation amplitudes than resting CD4^+^/CD8^+^ T cells in response to HyIL-6 treatment (Figure 1A and 1B). Interestingly, activated CD4^+^ T cells triggered higher levels of STAT1 phosphorylation than activated CD8^+^ T cells upon HyIL-6 stimulation (Figure 1A and 1B, left panels), which correlated with a higher expression of gp130 and IL-6Ra by CD4^+^ T cells (Sup. Figure 1). Overall, these results show that IL-6-induced signaling in human T cells is a very dynamic process.

**Figure 1.**
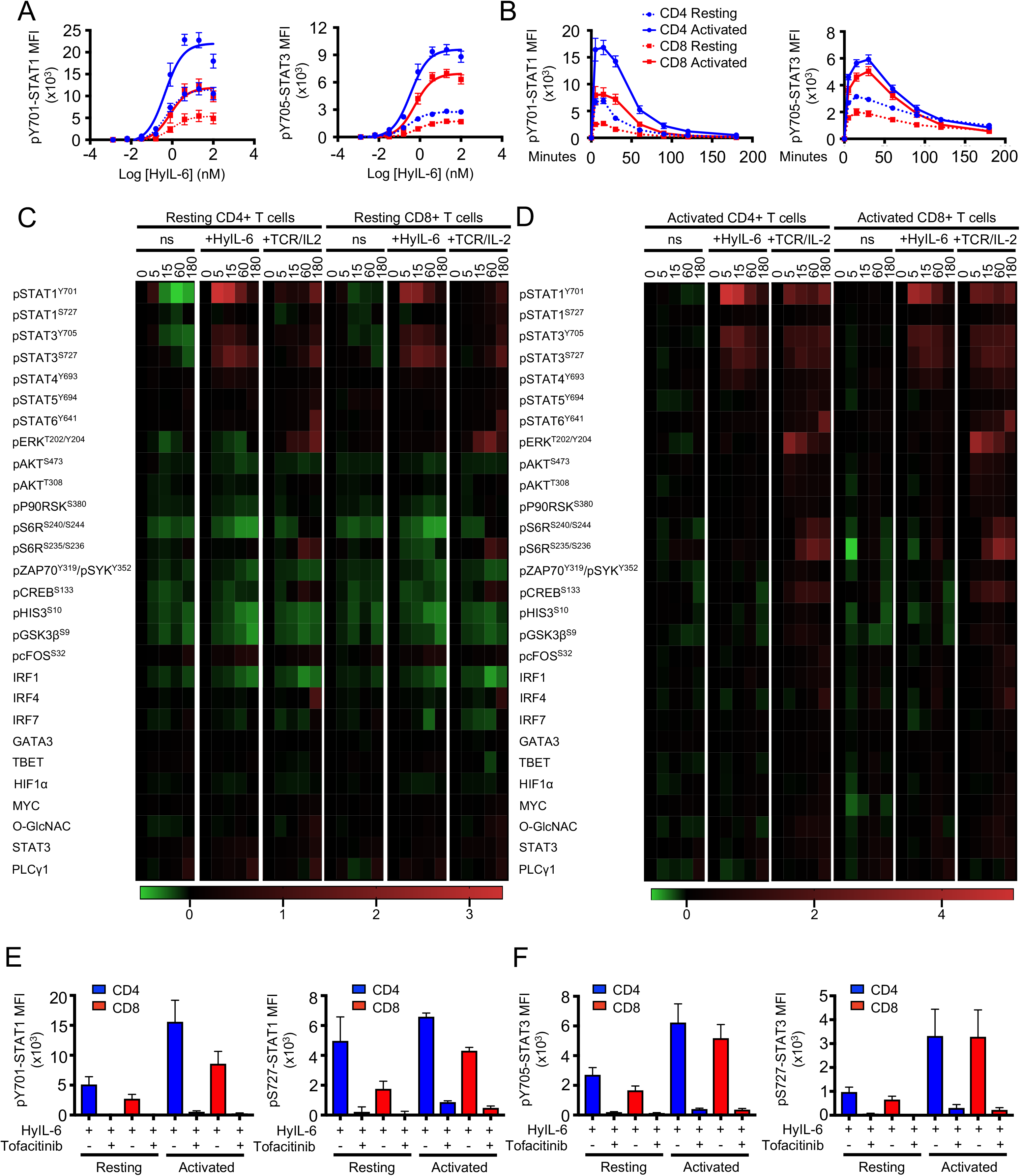
Inteleukin-6 signaling landscape in primary human T cells. A-B) STAT1 and STAT3 phosphorylation in response to varying doses (**A**) and exposure time (**B**) of Interleukin-6 stimulation in resting and activated primary human CD4^+^ and CD8^+^ T cells. **C-D)** Phospho-FLOW analysis of the main targets of IL-6 signaling pathway as described in the literature in resting (**C**) and activated primary human CD4^+^ and CD8^+^ T cells treated with HyIL-6 or anti-CD3/CD28 (TCR) + Interleukin-2. ns indicates cells without any stimulation. See also Supplementary Figure 2-3. **E-F)** Effect of JAK inhibition by 2μM Tofacitinib on the phosphorylation of STAT1 (**E**) and STAT3 (**F**) Tyr701 and Ser727 in resting and activated primary human CD4^+^ and CD8^+^ T cells. Error bars show mean + SEM.

To gain further insight into the signaling networks, beyond JAK/STAT1/3, engaged by IL-6 in T cells, we used an antibody array targeting 28 relevant signaling intermediaries. Resting and activated CD4^+^ and CD8^+^ T cells were stimulated with saturating concentrations of HyIL-6 for the indicated times and their signaling signatures in response to HyIL-6 treatment were assayed by flow cytometry (Figure 1C and 1D). To ensure the quality of our signaling antibody array, we stimulated the different populations of T cells with anti-CD3/anti-CD28 antibodies (TCR)+IL-2 as a positive control, since this treatment activates a large proportion of the signaling molecules detected by our antibody array (Ross et al., 2016; Smith-Garvin et al., 2009). In both resting and activated human CD4^+^ and CD8^+^ T cells, TCR+IL-2 treatment led to the activation of a vast array of signaling intermediaries, including STAT1, STAT3, STAT4, STAT5, STAT6, ERK, AKT, S6R, CREB (Figure 1C, 1D and Sup. Figure 2 and 3). HyIL-6 treatment on the other hand preferentially induced the tyrosine and serine phosphorylation of both STAT1 and STAT3, and to a lower extent STAT4 (Figure 1C, 1D and Sup. Figure 2 and 3). STAT1 and STAT3 Tyr and Ser phosphorylation in response to HyIL-6 treatment were inhibited by the JAK inhibitor Tofacitinib, confirming the dependency of these two modifications on JAK activity (Figure 1E, 1F and Sup. Figure 4 A-D). Overall our signaling data support a dynamic and exclusive activation of the JAK/STAT pathway by IL-6 in T cells, suggesting that IL-6 functional pleiotropy emanates from activation of a few signaling intermediaries.

### IL-6 induces large number of phosphoproteome changes in human T cells

To obtain a full spectrum of the IL-6 signalosome in T cells, we next performed quantitative high-resolution phosphoproteomics assay using stable isotope labeling by amino acids in cell (SILAC). We selected Th-1 cells due to their significance in immunity and ability to expand *in vitro* to large quantities of a highly pure population, compatible with SILAC studies. Briefly, CD4^+^ T cells were isolated from PBMCs of healthy donors using anti-CD4-FiTC (Biolegend) and anti-FITC microbeads (Miltenyi Biotec) and polarized into Th-1 for three days (Sekiya and Yoshimura, 2016). From day 3, cells were split into two independent cultures and expanded for additional ten days. Cells grown in light SILAC media (L-Arginine R0, L-Lysine K0) were used as unstimulated control and cells grown in medium SILAC media (L-Arginine U-^13^C_6_ R6, L-Lysine U-^13^C_6_ K6) were stimulated with HyIL-6 for fifteen minutes. Labelling incorporation greater than 97% was confirmed by Mass Spectrometry. Cells were mixed in a 1:1 ratio and lysed using 1% SDS buffer in 100mM Triethylammonium bicarbonate buffer (TEAB) and digested using MS-grade Trypsin (Thermo). Phosphopeptides were enriched using HILIC fractionation followed by MagResyn Ti-IMAC (ReSyn Biosciences) affinity chromatography and analysed on an Q Exactive^TM^ Plus Mass Spectrometer. Raw files were analyzed using MaxQuant (Figure 2A).

**Figure 2.**
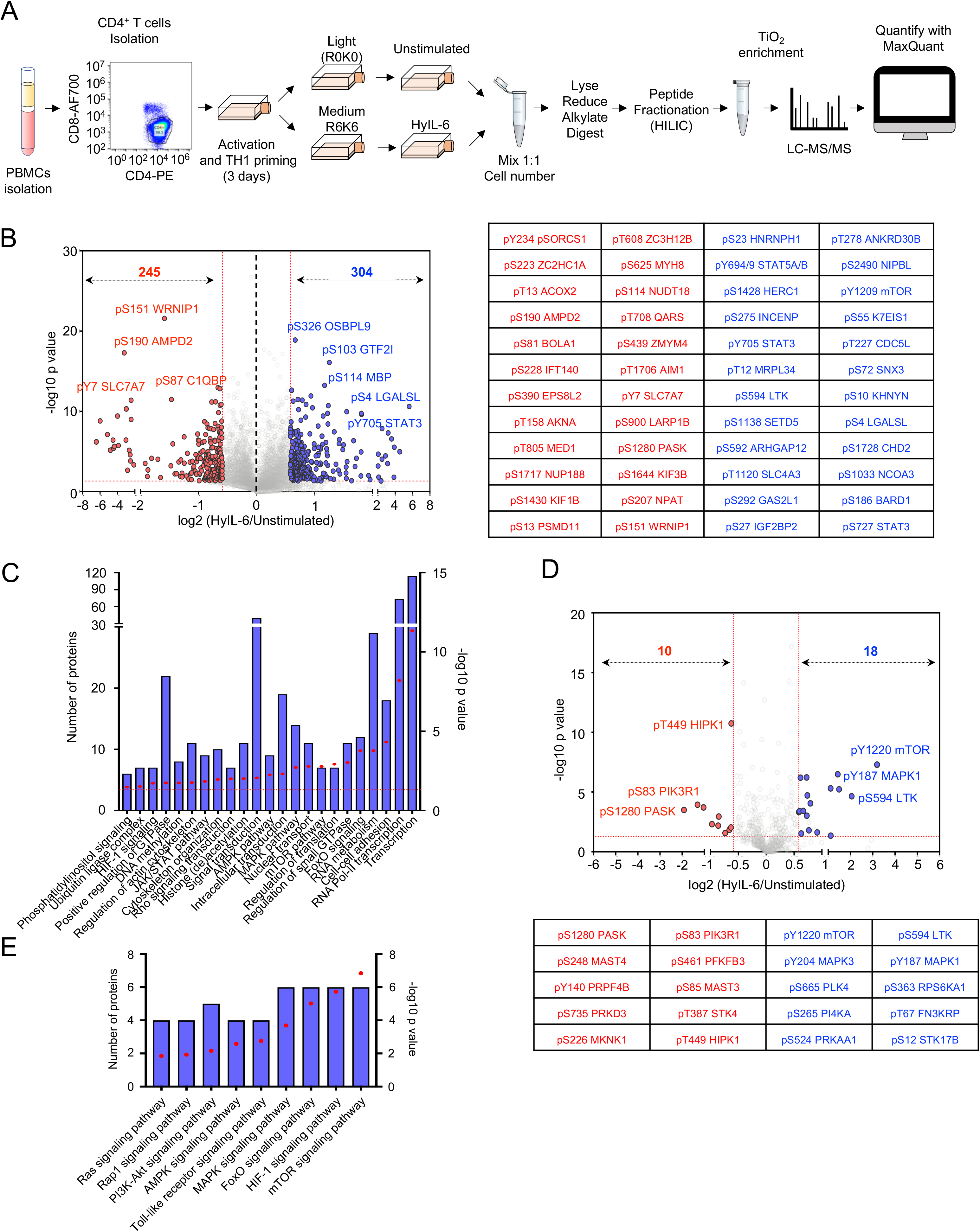
Phosphoproteomic landscape of Interleukin 6 in human primary CD4^+^ Th1 cells. **A)** Experimental workflow for SILAC-based quantitative phosphoproteomic analysis of human primary CD4^+^ Th1 cells stimulated with 20nM HyIL-6 for 15 minutes. **B)** Volcano plot showing differential phosphopeptides in unstimulated versus stimulated Th1 cells with 20nM HyIL-6 for 15 minutes. Phosphosites changed more than 1.5-fold with a p value <0.05 are shown in red (decreased) or blue (increased). Phosphopeptides identified in six biological replicates are shown as log-transformed SILAC ratios plotted against log-transformed p values (two-sided T test). Select phosphopeptides are labeled (see Supplementary Table S1 for full list). The 24 phosphosites more reproducibly decreased (red) or increased (blue) are displayed alongside. **C)** Gene ontology (GO) analysis as determined by DAVID of the phosphosites regulated by HyIL-6 in human primary Th1 cells. **D)** Kinases phosphosites regulated in response to HyIL6 stimulation in human primary Th1 cells. **E)** GO analysis as determined by DAVID showing main signaling pathways engaged by HyIL-6 in human primary Th1 cells.

The combined analysis from six independent biological replicates of unstimulated vs HyIL-6 stimulated Th-1 cells identified 17,935 phosphosites on 4,196 proteins (Figure 2B and Sup. Table S1). Among those 304 were increased and 245 were decreased in response to HyIL-6 stimulation in human primary Th-1 cells (Figure 2B). GO analysis (KEGG pathways and molecular function analyses) showed an enrichment of the JAK/STAT pathway as expected upon stimulation of Th-1 cells with HyIL-6 (Figure 2C). Our phosphoproteomic study identified 288 kinases (Figure 2D and Sup. Table S1). Of those 28 were regulated by HyIL-6 treatment (Figure 2D and Sup. Table S1) and were enriched in several signaling pathways (e.g. mTOR, MAPK, etc) (Figure 2E). Interestingly, nearly 40% of the phosphoproteomic changes induced by HyIL-6 took place in the nucleus (Figure 3 and Sup. Figure 5), highlighting this compartment as a major signaling platform for IL-6 activities. Furthermore, our GO analysis indicated an enrichment of phosphosites involved in the regulation of transcription and more specifically RNA pol II-mediated transcription (Figure 2C). HyIL-6 treatment regulated processes related to protein transcription (ELF2, GTF2I, TMF1), histone (de)acetylation (KANSL2, TRRAP, RBBP7), RNA pol II transcription (MECPE, MEF2C, MED1), mRNA splicing (DDX46, SF3B4), processing (RBM6, RBM39) and export (NUP50, NUP153) (Figure 3 and Sup. Figure 5A). Additionally, HyIL-6 treatment regulated non-nuclear processes, including regulation of the cytoskeleton, translation and proteasome at the cytoplasmic level (Figure 2C and Figure 3). Overall our phosphoproteomic revealed a strong regulation of the nuclear phosphoproteome by IL-6 that could contribute to fine-tune its immuno-modulatory activities.

**Figure 3.**
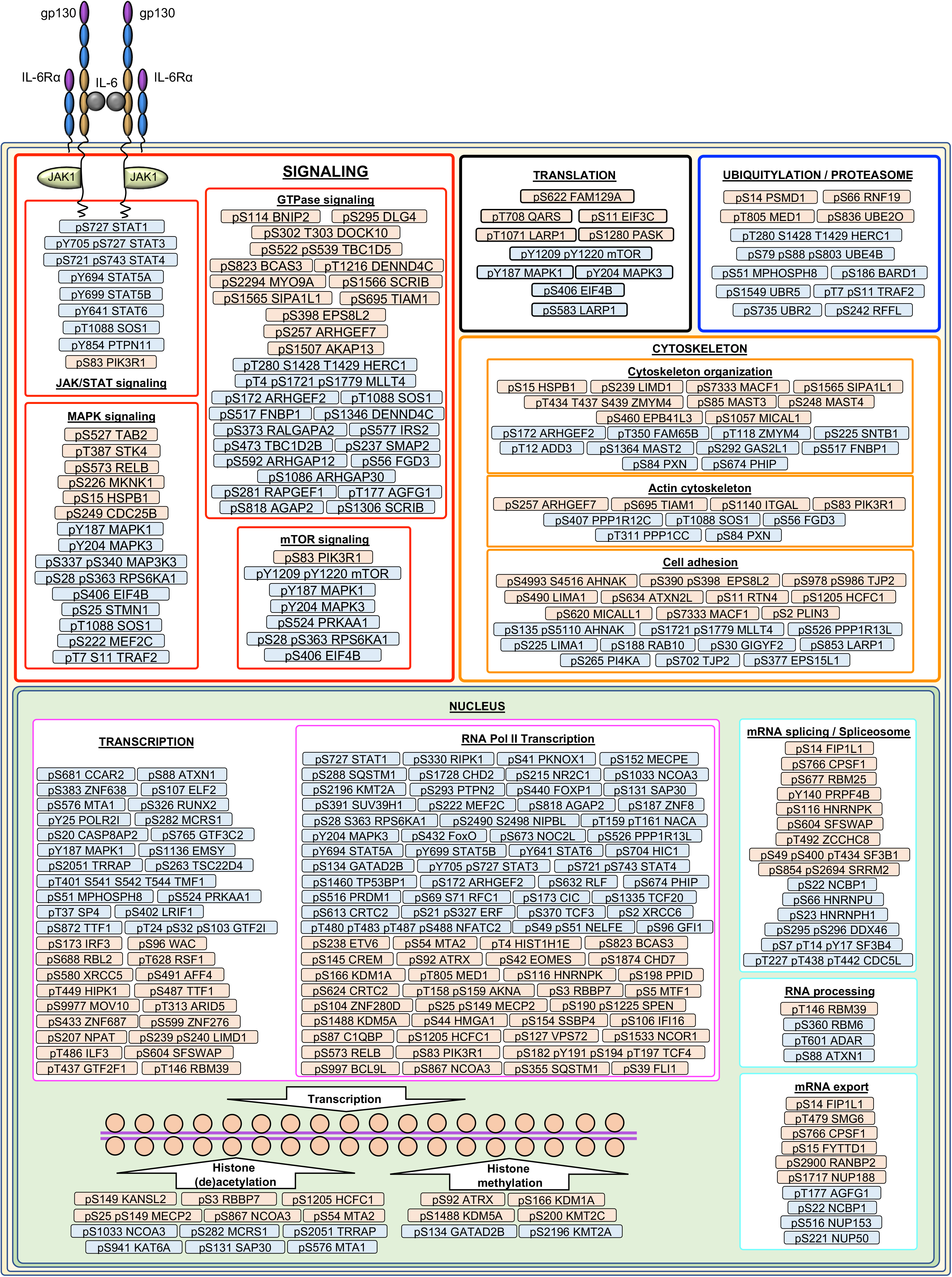
Proteins regulated by HyIL-6 in human primary CD4^+^ Th-1 cells. The scheme shows the cellular location (See Supplementary Figure 5) and molecular function of the proteins regulated by phosphorylation in response to HyIL-6 stimulation in human primary CD4^+^ Th-1 cells as determined by DAVID analysis. Refer also to Supplementary Figure 5.

### CDK8/CDK9 regulate STAT1 and STAT3 Ser727 phosphorylation

HyIL-6 triggers the Tyr and Ser phosphorylation of STAT1 and STAT3 in T cells. While JAK1 contributes to the Tyr phosphorylation of STAT1/3, the kinase responsible for STAT1/3 serine phosphorylation in T cells is currently not known. To identify this kinase, we used a panel of inhibitors targeting signaling pathways previously shown to regulate STAT1 and STAT3 serine phosphorylation in different cellular systems (Decker and Kovarik, 2000). Tofacitinib, a JAK inhibitor, blocked both STAT1/STAT3 Tyr and Ser phosphorylation by HyIL-6 treatment (Figure 4A and 4B) confirming previous observations. Of the battery of inhibitors tested, only Torin 1, an mTOR inhibitor targeting both mTORC1 and mTORC2 complexes (Thoreen et al., 2009), inhibited the Ser phosphorylation induced by HyIL-6 in both STAT1 and STAT3, without affecting their Tyr phosphorylation (Figure 4A and 4B). Rapamycin, an mTORC1-only complex inhibitor (Laplante and Sabatini, 2012) failed to do so, suggesting that the Ser phosphorylation of STAT1 and STAT3 induced by HyIL-6 was a mTORC2-mediated response (Figure 4A and 4B). However, alternative mTOR inhibitors (i.e. AZD8055 and KU0063794) failed to restrict STAT1 and STAT3 Ser phosphorylation by HyIL-6, indicating that the Torin1 mediated inhibition was an off-target effects (Figure 4C). Moreover, inhibitors specifically targeting well-described off-targets of Torin 1, i.e. Ataxia Telangiectasia Mutated (ATM; KU53933) and DNA-dependent Protein Kinase (DNA-PK; KU57788)(Liu et al., 2012), failed to inhibit STAT1 and STAT3 Ser phosphorylation by HyIL-6 (Figure 4D). This indicates that the inhibition of STAT1 and STA3 Ser phosphorylation is a previously unknown off-target effect of Torin1 and needs to be accounted for when used in *in vitro* or *in vivo* studies.

**Figure 4.**
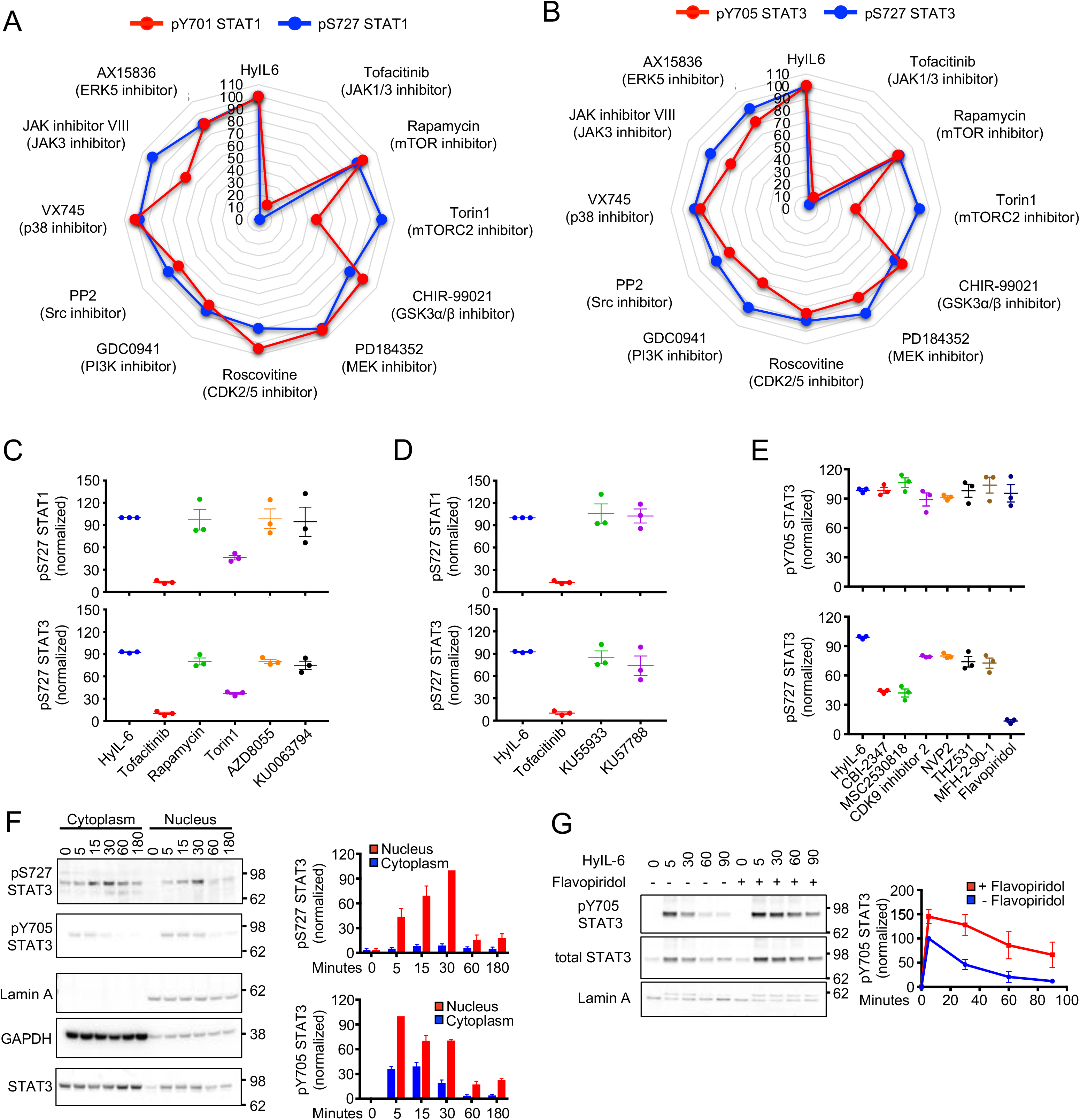
STAT1 and STAT3 HyIL-6-induced Ser727 phosphorylation is CDK8/9 mediated. A-B) Spider plots showing STAT1 (**A**) and STAT3 (**B**) pTyr701 (red line) and pSer727 (blue line) MFI normalized to HyIL-6-treated cells in the presence of different inhibitors in human primary CD4^+^ Th1 cells. **C)** Effect of different mTOR inhibitors on the STAT1 (upper panel) and STAT3 (lower panel) Ser727 phosphorylation induced by HyIL-6 in human primary CD4^+^ T cells. **D)** Effect of ATM inhibitor (KU53933) and DNA-PK inhibitor (KU57788) on the STAT1 (upper panel) and STAT3 (lower panel) Ser727 phosphorylation induced by HyIL-6 in human primary CD4^+^ T cells. **E)** Effect of different CDK inhibitors on the STAT3 Tyr705 (left panel) and STAT3 Ser727 (right panel) phosphorylation induced by HyIL-6 in human primary CD4^+^ T cells. **F)** Immunoblotting analysis of the cytoplasmic and nuclear fractions of human primary CD4^+^ Th1 cells stimulated with HyIL-6 for the times indicated. Quantitative analysis showing the combined data of three individual biological replicas are shown alongside. **G)** Time-course of STAT3 Tyr705 phosphorylation in nuclear fraction of human primary CD4^+^ cells stimulated with HyIL-6 in the absence (blue line) or presence (red line) of 2μM Flavopiridol. Quantitative analysis showing the normalized data of three independent biological samples is shown alongside. Error bars show mean + SEM.

Previous studies have described CDKs as regulators of STAT1 Ser phosphorylation in different systems (Bancerek et al., 2013; Chen et al., 2019; Kosciuczuk et al., 2019; Putz et al., 2013). Thus, we next tested whether the STAT3 Ser phosphorylation induced by HyIL-6 was mediated by CDKs. For that, we measured STAT3 Tyr and Ser phosphorylation levels induced by HyIL-6 in cells treated with a panel of CDK inhibitors. Flavopiridol, a well-described pan-CDK inhibitor (Luke et al., 2012), completely abolished the Ser phosphorylation of STAT3 induced by HyIL-6 (Figure 4E, Sup. Figure 6 A-C). CDK8 specific inhibitors (i.e. CBI-2347 and MSC2530818) inhibited HyIL-6 induced STAT3 Ser phosphorylation by 70% in both human CD4^+^ and CD8^+^ T cells (Figure 4E, Sup. Figure 6 A-C). CDK9 (i.e. NVP2 and CDK inhibitor II) or CDK12/CDK13 (i.e. THZ531 and MFH-2-90-1) inhibitors only reduced the STAT3 Ser phosphorylation by 20% (Figure 4E, Sup. Figure 6 A-C). None of the inhibitors affected the STAT3 Tyr phosphorylation levels (Figure 4E, Sup. Figure 6 A-C). These data highlight a critical role of CDK8 in regulating STAT3 Ser phosphorylation by HyIL-6, with an accessory role of other CDK members. In agreement with this, nuclear fractionation studies revealed that while STAT3 Tyr phosphorylation is fast and takes place in the cytoplasm, STAT3 Ser phosphorylation is delayed and occurs after STAT3 nuclear translocation (Figure 4F). Interestingly, inhibition of CDKs activity using Flavopiridol produced a prolonged STAT3 Tyr phosphorylation and the retention of Tyr phosphorylated and total STAT3 in the nuclear fraction (Figure 4G and Sup. Figure 6D). Similar observations were obtained for STAT1 (Sup. Figure 6E). Overall, our data suggest that CDKs control STAT3 nuclear dynamics by modulating its Ser phosphorylation levels.

### HyIL-6 induces nuclear interaction of STAT3 and CDK8/CDK9

Next, we explored whether STAT3 and CDK8/9 physically interacted in the nucleus upon HyIL-6 stimulation. For that, we performed Proximity Ligation Assays (PLA), a technique that allows the detection of protein complexes at endogenous levels without the need of protein overexpression or labelling that could interfere with their binding partners (Fredriksson et al., 2002). Activated human primary CD4^+^ T cells were stimulated with 20nM HyIL-6 for the indicated times and samples were prepared for PLA analysis following manufacturer instructions (Sigma). In untreated cells, we detected very low levels of STAT3/CDK8 (Figure 5A) and STAT3/CDK9 complexes (Figure 5B). Upon HyIL-6 stimulation, we detected a 2-4 fold increase in the number of STAT3/CDK8 and STAT3/CDK9 complexes, which peak at 30 min stimulation and return to basal levels after 3 hours (Figure 5A and 5B), paralleling the STAT3 Tyr activation kinetics (Figure 1B).

**Figure 5.**
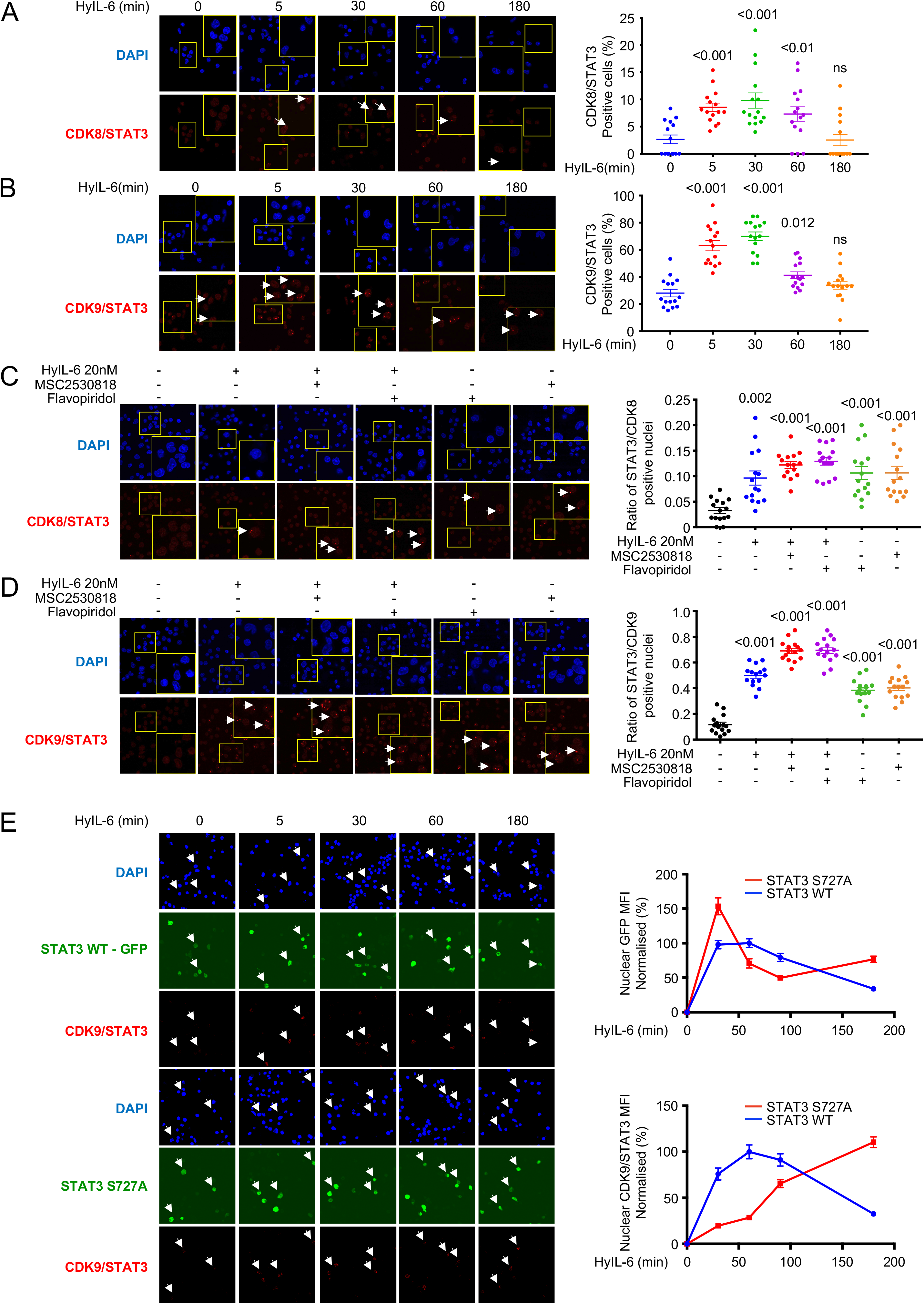
Proximity ligation assay analysis of the interaction of STAT3 and CDK8/9 induced upon HyIL-6 stimulation in human primary CD4^+^ Th1 cells. A-B) Kinetics of the STAT3/CDK8 (**A**) or STAT3/CDK9 (**B**) interaction induced by 20nM HyIL-6 in human primary CD4+ Th1 cells. **C-D)** STAT3/CDK8 (**C**) or STAT3/CDK9 (**D**) interactions were analyzed by PLA upon 20nM HyIL6 stimulation in the absence or presence of 2μM MSC2530818 or 2μM Flavopiridol, or upon treatment with the inhibitor only. White arrows in **A-D** indicate example of cells where interaction signal was detected. Cumulative plots from n=15 pictures alongside show the percentage of positive cells. P-values are calculated based on non-parametric twotailed Wilcoxon test against the control group (first bar on the left). **E)** STAT3/CDK9 interaction analyzed by PLA upon 20nM HyIL6 stimulation in STAT3 KO Hut78 cells expressing STAT3 WT-GFP (upper panel) or STAT3 Ser727Ala-GFP (lower panels). White arrows indicate example of cells expressing the recombinant protein and where the STAT3/CDK9 interaction was detected by PLA. Graph alongside shows the nuclear GFP MFI normalized to unstimulated cells (upper graph) or the nuclear STAT3/CDK9 PLA MFI in GFP positive cells normalized to unstimulated cells (lower graph). Quantitative data generated from n=15 pictures. Error bars show mean + SEM.

Our data showed that CDKs activity regulated STAT1/3 nuclear residency time (Figure 4G and Sup. Figure 6D and 6E). Thus, we next studied whether CDKs activity modulated the formation of STAT3/CDK complexes upon HyIL-6 stimulation. Activated human CD4^+^ T cells were stimulated with HyIL-6 for 30 min in the presence of different CDKs inhibitors. Levels of STAT3/CDK8 and STAT3/CDK9 complexes were measured by PLA analysis. As before, HyIL-6 stimulation led to a significant increase in the number of STAT3/CDK8 and STAT3/CDK9 complexes, when compared to unstimulated cells (Figure 5C and 5D). Addition of Flavopiridol (panCDK inh.) or MSC2530818 (CDK8 inh.) inhibitors resulted in an enhancement in the number of STAT3/CDK8 and STAT3/CDK9 complexes (Figure 5C and 5D). Interestingly, Flavopiridol (panCDK inh.) or MSC2530818 (CDK8 inh.) treatment alone led to an increase in the number of STAT3/CDK8 and STAT3/CDK9 complexes (Figure 5C and 5D), suggesting a role for CDKs activities in regulating STAT3 nuclear resident time and thus chromatin binding. Previous studies have reported that mutation of Ser727 in STAT3 to Ala modulates its transcriptional activity (Chung et al., 1997; Kim and Baumann, 1997; Lim and Cao, 1999; Wen et al., 1995; Yokogami et al., 2000). Thus, we next investigated the role that this mutation plays on recruitment of CDKs upon HyIL-6 stimulation. For that, we took advantage of the human Hut78 cell line, a cutaneous T lymphocyte, where HyIL-6 treatment also induced STAT3 Ser727 phosphorylation in a CDK dependent manner (Sup. Figure 7 A-C). Importantly, only Flavopiridol treatment resulted in an inhibition of STAT3 Ser phosphorylation by HyIL-6 in Hut78 cells, suggesting that in these cells CDK9 is the main CDK driving STAT3 phosphorylation (Sup. Figure 4A and 4B). Next, we generated STAT3 knock-down (STAT3 KnD) Hut78 cell lines by Crispr/CAS9 (Sup. Figure 7D). These cells exhibited a clear reduction in the STAT3 Tyr phosphorylation upon HyIL-6 stimulation (Sup. Figure 7E). STAT3 KnD cells were reconstituted with wt STAT3-GFP or Ser727A STAT3-GFP mutant and the levels of STAT3/CDK9 complex formation were measured by PLA (Figure 5E and Sup. Figure 7F). Due to STAT3 over-expression in these cells, we detected significantly higher levels of STAT3/CDK9 complex in unstimulated cells than those detected in human Th-1 cells. Yet, upon HyIL-6 stimulation we observed a significant increase in the number of STAT3 wt/CDK9 complexes in the nucleus, which peaked at 30 min and went back to basal levels by two hours treatment (Figure 5E). STAT3 Ser727A mutant exhibited a similar nuclear translocation profile to STAT3 wt, but it showed a delayed association kinetic with CDK9 (Figure 5E). Interestingly, at late stimulation times we observed higher levels of STAT3 Ser727Ala/CDK9 complexes when compared to STAT3 wt, suggesting that STAT3/CDK9 interaction is stabilized in the absence of STAT3 Ser phosphorylation (Figure 5E). Overall, our studies show that CDK8 and CDK9 fine-tune STAT3 nuclear dynamics via Ser phosphorylation.

### CDK8 regulates IL-6-induced nuclear phosphoproteome

Our data have highlighted a critical role of CDK8 in regulating STAT3 Ser phosphorylation and nuclear dynamics. Next, we asked which proportion of the IL-6 regulated phosphoproteome was dependent on CDK8 activity. For that, we performed phosphoproteomics studies in Th-1 cells stimulated with HyIL-6 for 15 min in the absence or presence of the CDK8 inhibitor MSC2530818. CD4^+^ T cells were isolated using anti-CD4-FiTC (Biolegend) and anti-FITC microbeads (Miltenyi Biotec) from PBMCs obtained from healthy donors and activated in Th-1 polarizing conditions for three days (Sekiya and Yoshimura, 2016). From day 3, cells were split into three independent cultures and expanded for ten days. Cells grown in light SILAC media (L-Arginine R0, L-Lysine K0) were used as unstimulated control, cells grown in medium SILAC media (L-Arginine U-^13^C_6_ R6, L-Lysine U-^13^C_6_ K6) were stimulated with HyIL-6 for fifteen minutes and cells grown in heavy SILAC media (L-Arginine U-^13^C_6_-^15^N_4_ R10, L-Lysine U-^13^C_6_-^15^N_2_ K8) were incubated with MSC2530818 2μM for 30 minutes prior to HyIL-6 stimulation for 15 minutes. Labelling incorporation greater than 97% was confirmed by Mass Spectrometry. Cells were mixed in a 1:1 ratio and lysed using 1% SDS buffer in 100mM Triethylammonium bicarbonate buffer (TEAB) and digested using MS-grade Trypsin (Thermo). Phosphopeptides were enriched using HILIC fractionation followed by MagResyn Ti-IMAC (ReSyn Biosciences) affinity chromatography and analyzed on an Q Exactive Plus Mass Spectrometer. Raw files were analyzed using MaxQuant (Sup. Figure 8).

The combined analysis of our phosphoproteomics study showed the identification of 11,035 phosphosites in 3,500 proteins (Figure 6A and 6B and Sup. table S2). HyIL-6 treatment induced the upregulation of 162 and downregulation of 160 phosphosites, where 88 of them (63 of the upregulated and 25 of the downregulated) were sensitive to CDK8 inhibition (Figure 6A and 6B and Sup. table S2). Consistently with our initial phosphoproteome study (Figure 2), most phosphoproteomic changes induced by HyIL-6 took place in the nuclei of the cells (Figure 6C). GO analysis studies indicated that HyIL-6 stimulation regulated proteins involved in transcription, specifically RNA pol II mediated transcription and other cellular processes such as histone (de)acetylation, and DNA methylation (Figure 6D and 6E). 27% of proteins involved in transcription and 40% of proteins involved in RNA Pol II transcription were affected by CDK8 inhibition (Figure 6F). A schematic view of the nuclear IL-6 induced phosphoproteome and its regulation by CDK8 is presented in Figure 6G. Overall, our data highlight a critical contribution of CDK8 in shaping the IL-6 phosphoproteome by regulating processes associated with RNA-Pol II mediated transcription.

**Figure 6.**
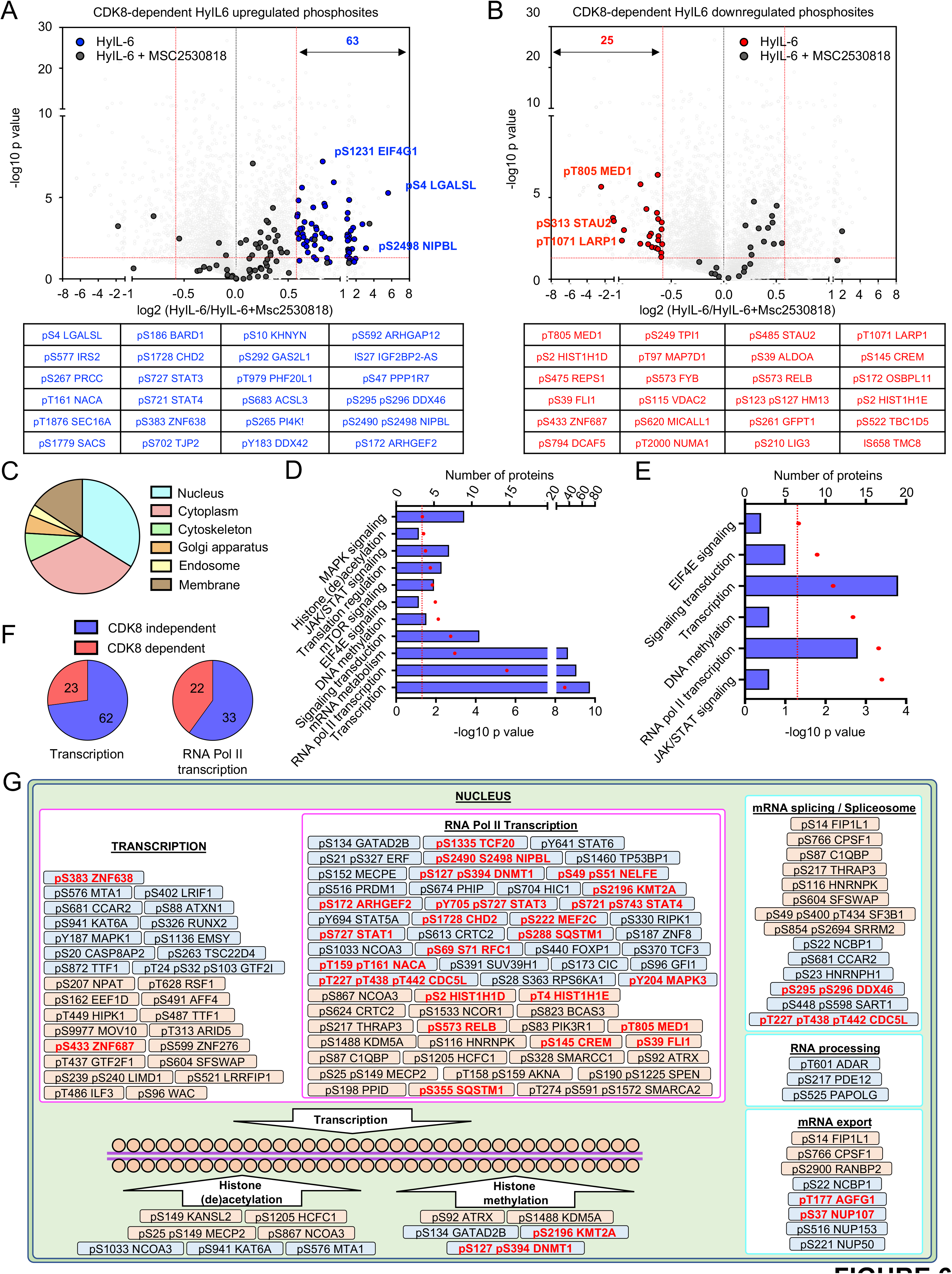
Regulation of the phosphoproteomic landscape of Interleukin 6 in human primary CD4^+^ Th-1 cells by CDK8. **A)** Volcano plot of the CDK8-dependent HyIL-6 upregulated phosphosites in human primary Th-1 cells (upper panel) and the 24 more affected phosphosites (lower panel). **B)** Volcano plot of the CDK8-dependent HyIL-6 downregulated phosphosites in human primary Th-1 cells (upper panel) and the 24 more affected phosphosites (lower panel). **C)** GO analysis showing the cellular location of the phosphosites regulated by HyIL-6 in a CDK8-dependent manner. **D)** GO analysis showing the main pathways and cellular processes regulated by HyIL-6 in human primary Th-1 cells. **E)** GO analysis showing the main pathways and cellular processes regulated by HyIL-6 in human primary Th-1 cells in a CDK8 dependent manner. **F)** Pie charts showing the number of phosphosites HyIL-6-regulated and CDK8-dependent phosphosites involved in the regulation of transcription (left graph) or Pol II-mediated transcription (right graph). **G)** The scheme shows the cellular location and molecular function of the proteins regulated by phosphorylation in response to HyIL-6 stimulation in human primary CD4^+^ Th-1 cells in a CDK8-dependent fashion as determined by DAVID analysis.

### CDK8 regulates STAT3 mediated transcription

We have shown that STAT3 nuclear dynamics were correlated with its Ser phosphorylation levels and were regulated by CDK8 activity. Thus, we next asked whether inhibition of CDK8 activity would alter STAT3-dependent gene transcription. For that, we performed RNA-Seq studies in Th-1 cells stimulated with HyIL-6 +/− the CDK8 inhibitor MSC2530818 for six hours. CDK8 inhibition by MSC2530818 did not alter the phosphorylation profile of RBP1, suggesting that our treatment did not affect basal transcriptional programs (Sup. Figure 9B) (Czodrowski et al., 2016; Harlen and Churchman, 2017). As previously observed by our laboratory (Martinez-Fabregas et al., 2019), HyIL-6 stimulation alone resulted in dysregulation of small number of genes (n=27) including classical STAT3 targets in Th-1 cells (Figure 7A and 7B-left panel). CDK8 inhibitor only treatment led to dysregulation of 111 genes of which 84 were up-regulated (Figure 7A and 7B-middle panel). The combined HyIL-6/MSC2530818 treatment exhibited a synergistic effect with 176 dysregulated genes upon treatment (Figure 7A and 7B-right panel), suggesting that CDK8 inhibition induced transcriptional programs by promoting HyIL-6/STAT3 mediated transcription. Dysregulated genes seem to fall into a few categories: genes induced by HyIL-6 treatment, but not regulated by CDK8 inhibition (e.g. BCL3 and SOCS3); genes induced upon CDK8 inhibition but not regulated by HyIL-6 treatment (e.g. AQP3 and CCR5); and genes exhibiting a synergic regulation by HyIL-6/CDK8 inhibition combined treatment (e.g. ACE and PDCD1) (Figure 7C). Next, we perform, gene set enrichment analysis for evaluating STAT3 mediated transcriptional activity in the absence and presence of CDK8 inhibitor. As expected, genes up-regulated by HyIL6 stimulation were highly enriched in genes known to be upregulated by STAT3 (GEO: GSE21670) (Figure 7D-top-panel). Importantly, however, genes up-regulated by CDK8 inhibition alone (Figure 7D-middle panel) or in combination with Hy-IL6 (Figure 7D-bottom panel) were also highly enriched in known STAT3 targets, indicating that Hy-IL6/STAT3 response is mediated via an intrinsic CDK8-dependent axis.

**Figure 7.**
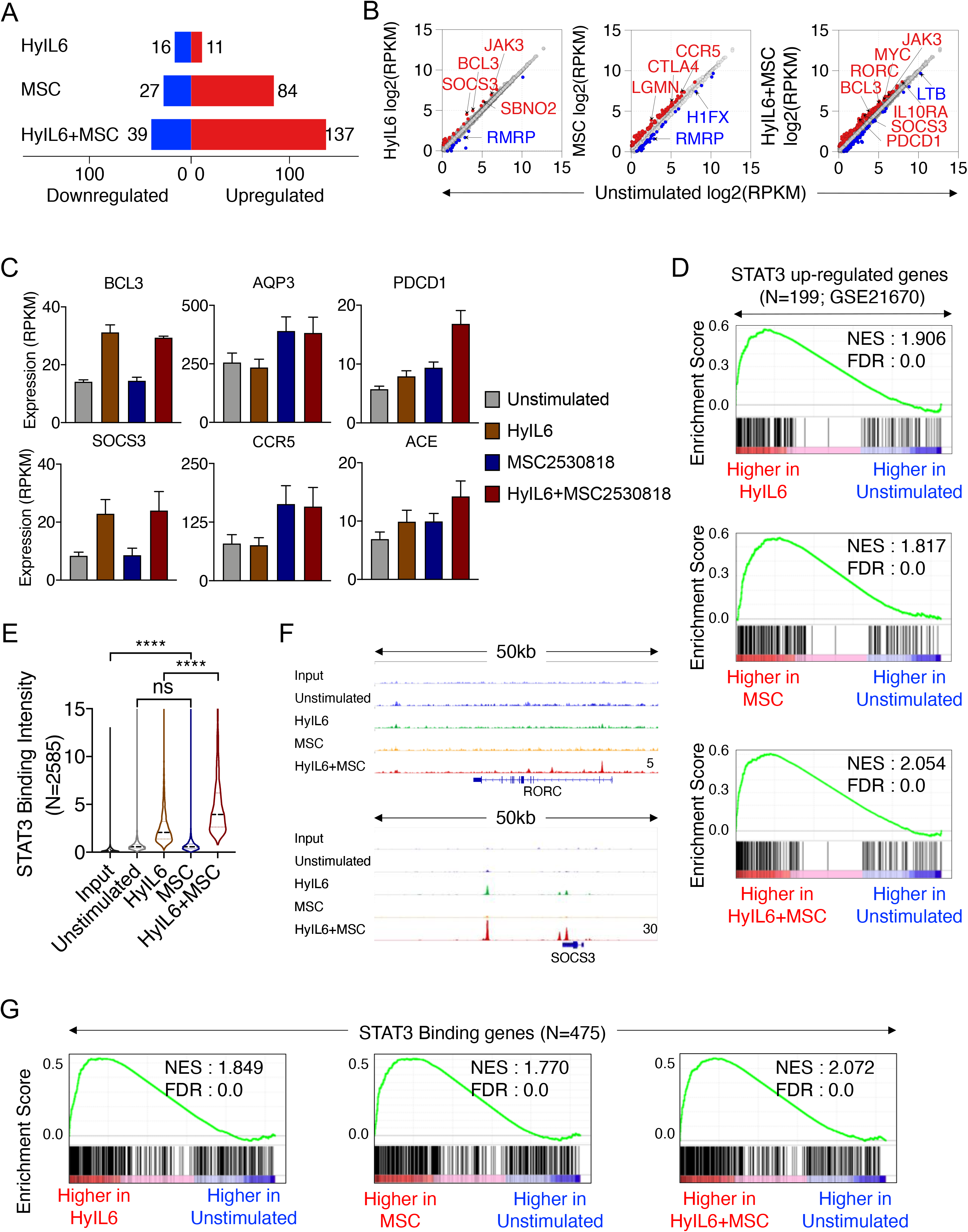
Transcriptional program elicited by interplay between HyIL-6 and CDK8 in human primary CD4^+^ Th1 cells. **A**) Number of differentially expressed genes (DEGs; fold change >1.5; p-value <0.05) between unstimulated versus HyIL6, MSC, or HyIL6+ MSC stimulated Th1 cells in n=3 donors. **B)** Scatter plot showing mean gene expression values (n=3) before (X-axis) and after indicated stimulation (Y-axis). Up-regulated (red) and downregulated (blue) genes are highlighted. **C**) Representative genes expression across different stimulation. Bars show mean + SEM. **D**) Geneset enrichment analysis (GSEA) (Subramanian et al., 2005) plots for STAT3 up-regulated genes (GSE21670) comparing stimulated versus unstimulated Th1 transcriptomes. NES: normalized enrichment score, FDR: false discovery rate. **E**) Violin plot showing the mean STAT3 binding intensity in n=2585 STAT3 bound regions across different stimulations. Peaks are identified by comparing Hy-IL6+MSC stimulation and input. P-values are determined by two-tailed Wilcoxon test (**** P<0.0001). **F**) Representative loci showing STAT3 binding across different stimulations. The height of the tracks are indicated at bottom corner of the plots. **G**) GSEA plots for 475 STAT3 bound genes comparing stimulated versus unstimulated Th1 transcriptomes.

Since CDK8 inhibition resulted in an accumulation of Tyr phosphorylated STAT3 in the nucleus, we next asked whether CDK8 could regulate STAT3 binding to chromatin in a genome-wide manner. To assess that, we carried out STAT3 ChIP-Seq in unstimulated or stimulated Th-1 cells with the three conditions described above for one hour. As expected, in unstimulated Th-1 cells, we detected very poor STAT3 DNA binding, which was significantly enhanced upon HyIL-6 treatment (Figure 7E). In Th-1 cells treated with the CDK8 inhibitor alone, we observed levels of STAT3 binding that resembled those obtained in unstimulated cells (Figure 7E). Strikingly, we observed a synergistic increase in binding intensity of STAT3 across target sites in cells stimulated with the combined HyIL-6/CDK8 inhibitor treatment when compared to HyIL-6 treatment alone, suggesting that inhibition of CDK8 activity amplify HyIL-6-induced STAT3 binding intensity to its target sites (Figure 7E with representative loci 7F). Importantly, in all conditions tested, STAT3 was binding to a canonical STAT3 GAS sequence motif (Sup. Figure 9C). Moreover, genes that were upregulated upon stimulation, were highly enriched in genes that harbor STAT3-binding site in their vicinity (Figure 7G), reaffirming a specific effect of CDK8 inhibition on STAT3 transcriptional activity. Collectively, our data confirm that CDK8 fine-tunes STAT3 binding dwell-time to chromatin via its Ser phosphorylation.

### CDK8 regulates Th-17 differentiation

IL-6 regulates inflammatory processes by inducing the differentiation of Th-17 cells. Moreover, we observed an increased binding of STAT3 at RORC gene locus upon stimulation with HyIL6 and CDK8 inhibitor (Figure 7F). Thus, we asked whether CDK8 activity would modulate Th-17 differentiation. Human primary resting CD4^+^ T cells were isolated from buffy coats and primed for 5 days under Th-17 polarizing conditions (Sekiya and Yoshimura, 2016). Primed cells were further expanded in media containing IL-2, anti-IL-4, anti-IFNγ in the presence or absence of the CDK8 inhibitor MSC2530818 (Figure 8A). At day 5 or 10 of expansion cells were analyzed by flow cytometry and ELISA for expression of the indicated cytokines and transcription factors (Figure 8). As previously described, HyIL-6 treatment induced a minor increase in the number of human Th-17 cells, highlighting the difficulty of working with these cells *in vitro* (Hakemi et al., 2011; Miyahara et al., 2008). About 2-3% of the T cells at the end of the polarization protocol were positive for IL-17 (Figure 8B and 8D) and RORgt expression (Figure 8F and 8H). CDK8 inhibition led to an average of a 3-fold increase in the number of IL-17 positive cells after 10 days expansion (Figure 8C and 8D). Importantly, we did not see changes in RORgt, IFNg or IL-4 expression by the T cells stimulated in the presence of the CDK8 inhibitor (Figure 8E-I) suggesting a specific regulation of STAT3-mediated Th-17 differentiation by the CDK8 inhibitor. Moreover, CDK8 inhibition resulted in an increase in the levels of secreted IL-17 as measured by ELISA (Figure 8J), but not IFNg (Figure 8K). Overall, our data agree with a model where CDK8 modulates STAT3 transcriptional processivity by fine-tuning its gene loci binding dwell-time, leading to a negative regulation of IL-6-mediated activities.

**Figure 8.**
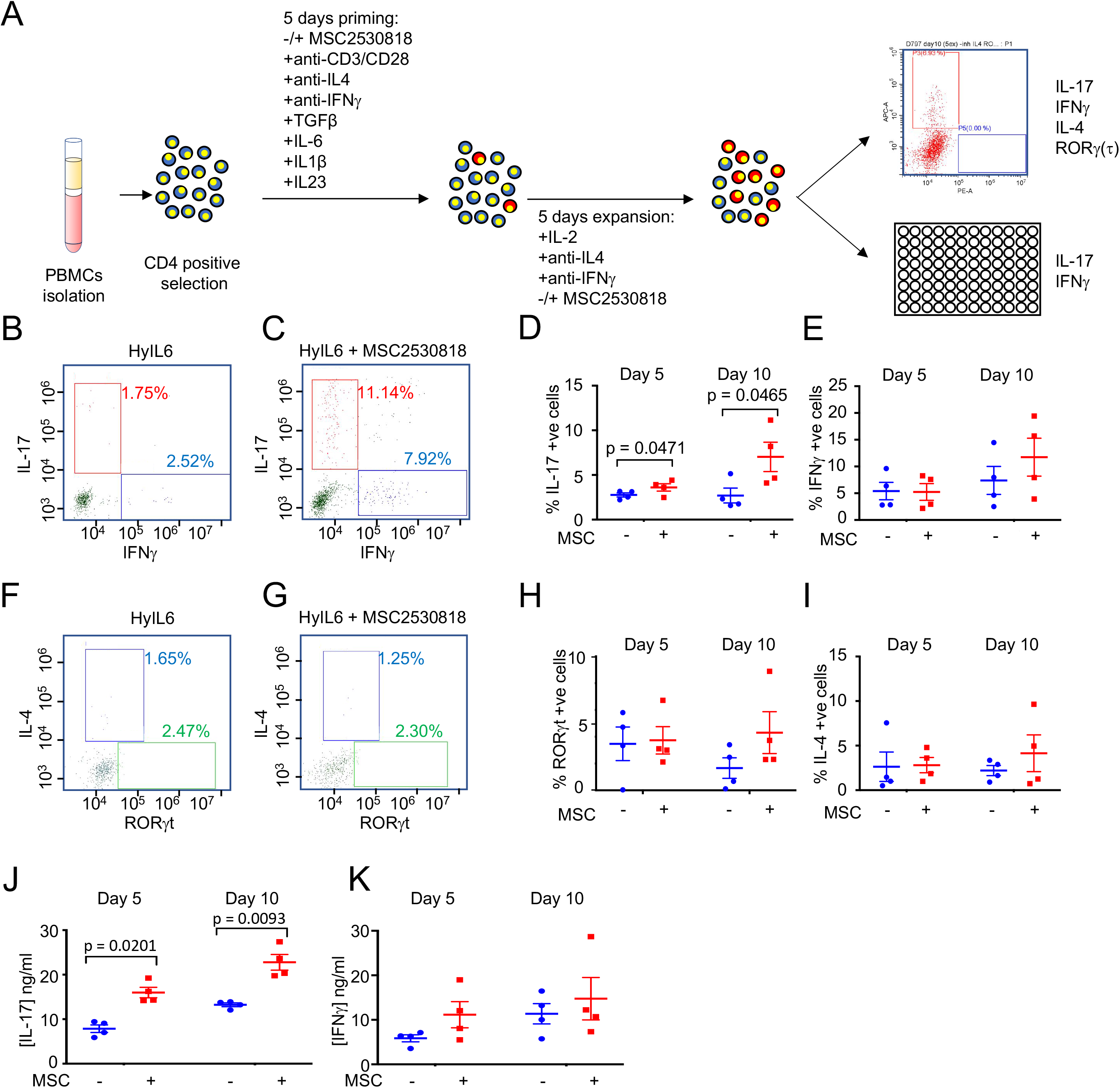
Role of CDK8 Ser727 phosphorylation of STAT3 in Th-17 differentiation in vitro. **A)** Experimental workflow for human Th-17 differentiation in vitro from isolated human resting CD4^+^ T cells. **B)** Dot plot representations of IL-17 and IFNγ positive cells in populations grown in the presence of HyIL-6. **C)** Dot plot representations of IL-17 and IFNγ positive cells in populations grown in the presence of HyIL-6 and MSC2530818. **D)** IL-17 positive cells were identified by flow-cytometry in untreated cells or cells treated with 2μM CDK8 inhibitor MSC2530818. Data are percentage of positive cells ± SEM in four donors; p values were calculated using a paired t-test. **E)** As in **D)** but for IFNγ positive cells. **F)** Dot plot representations of IL-4 and RORγt positive cells in populations grown in the presence of HyIL-6. **G)** Dot plot representations of IL-4 and RORγt positive cells in populations grown in the presence of HyIL-6 and MSC2530818. **H)** RORγt positive cells were identified by flow-cytometry in untreated cells or cells treated with 2μM CDK8 inhibitor MSC2530818. Data are percentage of positive cells ± SEM in four donors; p values were calculated using a paired t-test. **I)** As in **H)** but for IL-4 positive cells. **J)** Amount of IL-17 detected in growth media following growth of cells minus or plus inhibitor. **K)** Amount of IFNγ detected in growth media following growth of cells minus or plus inhibitor.

## DISCUSSION

Cytokines are highly pleiotropic ligands that act as master regulators of the immune response. Here, we have used IL-6 as a model system to uncover the molecular bases driving cytokine functional pleiotropy. We have identified CDK8 as a master regulator of the IL-6-induced signalosome and STAT3-dependent transcriptional activities. CDK8 is recruited to STAT3 binding gene loci upon IL-6 stimulation, where it triggers STAT3 Ser phosphorylation, and limits STAT3 chromatin resident time. Additionally, CDK8 regulates a large percentage of IL-6-induced nuclear phosphoproteome, further modulating IL-6 mediated transcription. This work outlines a general strategy to fine-tune cytokine responses by targeting CDKs activities to improve STAT transcriptional output.

Surprisingly, we found that nearly forty percent of the phosphosites induced by IL-6 treatment belonged to nuclear proteins, highlighting the key role for this compartment in defining IL-6 responses. These phosphosites were in the majority Ser/Thr phosphorylation events suggesting that JAKs were not the kinases involved in their regulation. Additionally, a closer look to the IL-6-induced kinome did not reveal an obvious signaling pathway induced by IL-6 that would account for its nuclear regulation. Thus, how does IL-6 treatment regulates the nuclear phosphoproteome? Our data showed that STAT3 binds CDK8 and CDK9 in the nucleus, where these two kinases drive its Ser phosphorylation. Inhibition of CDK8 activity resulted in the loss of a significant percentage of the IL-6-induced nuclear phosphoproteome. However, we did not detect phosphorylation of these two kinases by IL-6 treatment, suggesting that IL-6 might regulate their activity by altering their localization. Interestingly, several studies have reported that STAT3 can engage different gene expression programs in a cell-dependent and stimulating condition-dependent manner (Hutchins et al., 2013). These observations suggest a model where interaction between nuclear STAT3 and different kinases and/or transcription factors ultimately defines the STAT3 transcriptional program. In agreement with this model, several studies have described the interaction of STAT3 with different transcriptional factors in the promoter of responsive genes (Hutchins et al., 2013). In addition to changes in the nuclear phosphoproteome, our phosphoproteome study also identified phosphosites changes induced by IL-6 in proteins localized in endo-membranes (Golgi Apparatus, Endosomes, Lysosome, etc), cytoskeleton and cytoplasm, all compartments where STAT3 plays a critical role (Debidda et al., 2005; Jonchere et al., 2013; Liu et al., 2018; Lloyd-Lewis et al., 2018; Martinez-Fabregas et al., 2018; Ng et al., 2006; Shah et al., 2006; Teng et al., 2009). These phosphosites were also enriched in Ser/Thr phosphorylation. It is thus tempting to speculate, that in addition to its well-known transcriptional activity, which it is driven by a Tyr phosphorylation event via JAK activation, STAT3 acts as a signaling hub recruiting different Ser/Thr kinases in different cell compartments to fine tune cellular responses. This could explain why unlike other STAT members STAT3 gene inactivation is embryonic lethal (Takeda et al., 1997). Future studies will be required to fully characterize STAT3-mediated signaling complexes.

The role that CDK8 plays in transcription regulation has been controversial. While initially CDK8 was identified as a negative regulator of Pol-II mediated transcription (Jeronimo and Robert, 2017; Knuesel et al., 2009), more recent studies have described that CDK8 can also act as a positive modulator of Pol-II transcriptional activities (Chen et al., 2017; Donner et al., 2010; Donner et al., 2007). Our results suggest a negative role of CDK8 in STAT3 mediated transcription. CDK8 Ser phosphorylates STAT3 triggering its dissociation from chromatin and terminating STAT3-mediated transcription. Supporting our model, a recent report described that STAT3 Ser phosphorylation destabilizes the STAT3 Tyr phosphorylated homo-dimer, leading to its dissociation from chromatin (Yang et al., 2020). Interestingly, a series of recent studies have described that modulation of CDK8 activity, via either small-molecule inhibitors or genetic deletion, fine-tune responses elicited by different immune cells. Specific deletion of CDK8 in NK cells results in an enhancement of their anti-tumor responses (Witalisz-Siepracka et al., 2018), an activity that heavily relies in the action of different cytokines on NK cells (Hu et al., 2019). Moreover, small molecule inhibitors targeting CDK8 promote the differentiation of T regulatory (reg) cells, and Th-17 cells when T cells were placed under T reg polarizing conditions (Akamatsu et al., 2019; Guo et al., 2019), or Th-17 polarizing conditions (Figure 8) respectively. In both instances the polarizing conditions were enriched in different cytokines, highlighting potentially a broad regulation of cytokine responses by CDK8. Interestingly, CDK8 expression is a very dynamic process and it is often expressed at very low levels in cells. Naïve T cells express non-detectable levels of CDK8, which are upregulated upon T cells activation and differentiation (Howden et al., 2019). These observations suggest that cells can regulate their CDK8 levels to establish different thresholds of cytokine sensitivity. In agreement with this model, we detected a poor IL-6 induced gene expression program in Th-1 cells, which express high levels of CDK8 (Howden et al., 2019), despite high levels of STAT3 Tyr phosphorylation and STAT3 chromatin binding induced by IL-6 treatment.

Previous reports have highlighted a positive role of STAT Ser phosphorylation in STAT-mediated transcriptional activities. STAT1 Ser727Ala Knock-in mice elicited a poor immune response against *Listeria monocytogenes* infection (Varinou et al., 2003). Moreover, several studies have shown that Ser727Ala mutation impairs transcriptional activity by different STATs (Levy and Darnell, 2002; Lim and Cao, 1999; Sadzak et al., 2008; Wen et al., 1995). Yet our study and others (Chung et al., 1997; Decker and Kovarik, 2000; Kim and Baumann, 1997) highlight a negative role of STAT3 Ser phosphorylation in STAT3 nuclear dynamics and transcriptional activities. An important difference between our approach and previous studies using STAT Ser mutants is that in the latter the activity of CDK8 remained intact. We showed that CDK8 inhibition increases STAT3 chromatin binding and induces an upregulation of STAT3-dependent gene expression, establishing CDK8 as a negative regulator of STAT3-mediated transcription. Moreover, we could detect CDK8-mediated phosphorylation of different transcription factors in an IL-6-dependent manner (e.g. CREM, RelB and FLI1), indicating that CDK8 elicits a broader regulation of transcription machinery. Additionally, we have shown that STAT3 Ser727Ala mutation stabilizes the CDK/STAT3 interaction upon IL-6 stimulation. All together these observations suggest that the longer-lived CDK8/STAT3 complex formed in the context of the STAT3 Ser mutant, brings signaling active CDK8 to STAT3-dependent genes, resulting in transcriptional repression. Overall, our study provides new molecular evidences that establish CDK8 as a master regulator of STAT3 transcriptional activities and present a new strategy to harness the therapeutic potential of cytokines by finetuning CDK8 expression levels and activities in different immune cells.

## ACKNOWLEDGMENTS

We thank members of the Moraga, Mitra, and Majid laboratories for helpful advice and discussion. This work was supported by the Wellcome-Trust-202323/Z/16/Z and ERC-206-STG grant (I.M.), and National Heart, Lung and Blood Institute (K22HL125593, MK).

## COMPETING INTERESTS

The authors declare that they have no competing interests.

## MATERIALS AND METHODS

### Protein expression and purification

HyIL-6 (Fischer et al., 1997) cloned into the pAcGP67-A vector (BD Biosciences) in frame with an N-terminal gp67 signal sequence and a C-terminal hexahistine tag was produced using the baculoviral expression system, as previously described (LaPorte et al., 2008). The baculoviral stocks were prepared in *Spodoptera frugiperda (Sf9)* cells grown in SF900III media (Invitrogen, #12658027) and used to infect *Trichoplusiani ni* (High Five) cells grown in InsectXpress media (Lonza, #BELN12-730Q) for protein expression. After 48h infection, secreted protein was captured from High Five supernatants using HisPur Ni-NTA resin (Thermo Scientific, #88223) affinity chromatography, concentrated, and purified by size exclusion chromatography on a Enrich SEC 650 1 x 300 column (Biorad), equilibrated in 10 mM HEPES (pH 7.2) containing 150 mM NaCl. Recombinant HyIL-6 was purified to greater than 98% homogeneity.

### CD4^+^ T cell isolation

Peripheral blood mononuclear cells (PBMCs) from healthy donors were purified from buffy coats (Scottish Blood Transfusion Service) by density gradient centrifugation following manufacturer’s instructions (Lymphoprep, StemCell Technologies, #07801). For CD4^+^ T cells isolation, 1 x 10^8^ PBMCs per donor were stained with anti-CD4^FiTC^ antibodies (Biolegend, #357406) and isolated by magnetic activated cell sorting (MACS, Miltenyi) using anti-FiTC microbeads (Miltenyi, #130-048-701) according to manufacturer’s instructions to a purity ≥99%.

### Dose-response and kinetic experiments

For dose-response experiments of STAT1 or STAT3 phosphorylation, 96-well plates were prepared with 30μL of cells at 5 x 10^5^ cells/mL. Cell were then stimulated with different concentrations to obtain the dose-response curves. After stimulation cells were fixed with 2% formaldehyde for 10 minutes at RT.

For kinetics experiments, cell suspensions were stimulated with saturating concentrations of the cytokines (10 nM HyIL-6) as indicated and cells finally fixed with 2% formaldehyde for 10 minutes at RT.

### Permeabilization, fluorescence barcoding and antibody staining

After fixation, cells were collected by centrifugation at 1200 rpm for 5 min, formaldehyde blocked by washing the cells with 200 μL of PBS containing 5 mg/ml BSA (PBSA) and collected again by centrifugation at 1200 rpm for 5 min. Then, cells were resuspended and permeabilized in ice-cold methanol for 20 minutes on ice. Cells were then fluorescently barcoded (Krutzik and Nolan, 2006) using a combination of different concentrations of aminoacid reactive dyes (PacificBlue #10163, DyLight800 #46421, Thermo Scientific). Finally, cells were pooled and stained with anti-CD3^BV510^ (Biolegend, #300448), anti-CD4^PE^ (Biolegend, #357404), anti-CD8^AF700^ (Biolegend, #300920), anti-pSTAT1-Tyr701^AF647^ (Cell Signaling, #8009S), anti-pSTAT1-Ser727^AF488^ (Biolegend, #686410), anti-pSTAT3-Tyr705^AF488^ (Biolegend, #651006) and anti-pSTAT3-Ser727^AF647^ (Biolegend, #698914). Cells were analysed in a CytoFlex S flow cytometer (Beckman Coulter) with the individual cell populations being identified by their barcoding pattern and mean fluorescence intensity (MFI) for the different forms of STAT 1 or STAT3 measured.

### Phospho-FLOW

Resting PBMCs isolated as described before from buffy coats or upon activation for three days with ImmunoCult Human CD3/CD28 T Cell Activator (StemCell, #10971) following manufacturer instructions in the presence of 20 ng/mL IL2 (Novartis, #709421) were starved for 24 hours in RPMI 1640 (Invitrogen) containing 10% fetal bovine serum (FBS, Invitrogen, #A3160801) and then stimulated with 10nM HyIL-6 or 0.1 μg/mL anti-CD3 (Biolegend #300438) and 20 ng/mL IL2 (Novartis #709421). Then, cells were fixed with 2% formaldehyde, permeabilized with ice-cold methanol and barcoded as described above. Finally, cells were pooled and stained with anti-CD3^BV510^ (Biolegend, #300448), anti-CD4^PE^ (Biolegend, #357404), anti-CD8^AF700^ (Biolegend, #300920), anti-pSTAT1-Y701^AF647^ (Cell Signaling, #8009S), anti-pSTAT1-SER727ALA^F488^ (Biolegend, #686410), anti-pSTAT3-Y705^AF488^ (Biolegend, #651006) and anti-pSTAT3-SER727ALA^F647^ (Biolegend, #698913), anti-pSTAT4-Y693^AF488^ (BD Biosciences, #558136), anti-pSTAT5-Y694^AF647^ (Cell Signaling, #9365S), anti-pSTAT6-Y641 (BD Biosciences, #612600), anti-pERK-T202/Y204^AF488^ (eBiosciences, #53-9109-41), anti-pAKT-S473^AF488^ (Cell Signaling, #4071S), anti-pAKT-T308^AF647^ (Cell Signaling, #48646S), anti-pP90RSK-S380^AF488^ (Cell Signaling, #13588S), anti-pS6R-S240/S244^AF488^ (Cell Signaling, #5018S), anti-pS6R-S235/S236^AF647^ (Cell Signaling, #4851S), anti-pZAP70-Y319/pSYK-Y352^AF647^ (Cell Signaling, #82975S), anti-pCREB-S133^AF488^ (Cell Signaling, #9187S), anti-pHIS3-S10^AF647^ (Cell Signaling, #9716S), anti-pGSK3β-S9 (Cell Signaling, #14332S),anti-pCFOS-S32^AF647^ (Cell Signaling, #8677S), anti-IRF1^AF647^ (Cell Signaling, #14105S), anti-IRF4^AF647^ (Biolegend, #646408), anti-IRF7^AF647^ (Biolegend, #656007), anti-GATA3^AF488^ (Biolegend, #653807), anti-TBET^AF647^ (Biolegend, #644803), anti-HIF1α^AF488^ (Biolegend, #359707), anti-MYC^AF488^ (Cell Signaling, #12855S), anti-O-GlcNAC^AF647^ (NOVUS Biologicals, #NB300-524AF647), anti-STAT3A^PC^ (BD Biosciences, #560392) and anti-PLCγ1^AF647^ (BD Biosciences, #557883). Cells were analyzed in a CytoFlex S flow cytometer (Beckman Coulter) with the individual cell populations being identified by their barcoding pattern and mean fluorescence intensity (MFI) measured.

### Phosphoproteomics

Resting CD4^+^ T cells were labeled with anti-CD4-FiTC antibody (Biolegend, #357406) and isolated from human PBMCs by magnetic activated cell sorting (MACS, Miltenyi) using anti-FiTC microbeads (Miltenyi, Cat#130-048-701) following manufacturer instructions. Subsequently, resting CD4^+^ T cells were activated under Th-1 polarizing conditions. Briefly, 3×10^7^ resting human CD4^+^ T cells per donor were primed for three days with ImmunoCult Human CD3/CD28 T Cell Activator (StemCell, #10971) following manufacturer instructions in the presence of 20 ng/mL IL2 (Novartis #709421), 20 ng/mL IL12 (BioLegend, #573002) and 10 ng/mL anti-IL4 (BD Biosciences, #554481). Then, cells were split into three different conditions light SILAC media (40 mg/mL L-Lysine K0 (Sigma, #L8662) and 84mg/mL L-Arginine R0 (Sigma, #A8094)), medium SILAC media (49 mg/mL L-Lysine U-13C6 K6 (CKGAS, #CLM-2247-0.25) and 103 mg/mL L-Arginine U-13C6 R6 (CKGAS, #CLM-2265-0.25)) and heavy SILAC media (49.7 mg/mL L-Lysine U-13C6,U-15N2 K8 (CKGAS, #CNLM-291-H-0.25) and 105.8 mg/mL L-Arginine U-13C6,U-15N2 R10 (CKGAS, #CNLM-539-H-0.25)) prepared in RPMI SILAC media (Thermo Scientific, #88365) supplemented with 10% dialyzed FBS (HyClone, #SH30079.03), 5 mL L-Glutamine (Invitrogen, #25030024), 5 mL Pen/Strep (Invitrogen, #15140122), 5 ml MEM vitamin solution (Thermo Scientific, #11120052), 5 mL Selenium-Transferrin-Insulin (Thermo Scientific, #41400045) and expanded in the presence of 20 ng/mL IL2 and 10 ng/ml anti-IL4 for another 10 days in order to achieve complete labelling. Incorporation of medium and heavy version of Lysine and Arginine was checked by mass spectrometry and samples with an incorporation greater than 95% were used. After expansion, cells were starved without IL2 for 24 hours before stimulation with 10 nM HyIL-6 in the presence or absence of 2 μM MSC2530818 (Stratech, # S8387-SEL) for 15 minutes. Cells were then washed three times in ice-cold PBS, mix in a 1:1:1 ratio, resuspended in SDS-containing lysis buffer (1% SDS in 100mM Triethylammonium Bicarbonate buffer (TEAB)) and incubated on ice for 10 minutes to ensure cell lysis. Then, cell lysates were centrifuged at 20000 g for 10 minutes at +4°C and supernatant was transferred to a clean tube. Protein concentration was determined by using BCA Protein Assay Kit (Thermo, #23227), and 10 mg of protein per experiment were reduced with 10mM dithiothreitol (DTT, Sigma, #D0632) for 1 hour at 55°C and alkylated with 20mM iodoacetamide (IAA, Sigma, #I6125) for 30 min at RT. Protein was then precipitated using six volumes of chilled (−20°C) acetone overnight. After precipitation, protein pellet was resuspended in 1mL of 100mM TEAB and digested with Trypsin (1:100 w/w, Thermo, #90058) and digested overnight at 37°C. Then, samples were cleared by centrifugation at 20000 g for 30 min at +4°C, and peptide concentration was quantified with Quantitative Colorimetric Peptide Assay (Thermo, #23275). Phosphopeptide enrichment in the peptide fractions generated as described above was carried out using MagResyn Ti-IMAC following manufacturer instructions (2BScientific, MR-TIM002). Phosphopeptide samples were analyzed using a nanoflow liquid chromatography system (Ultimate 3000 RSLCnano system, Thermo Scientific) coupled to a Q Exactive Plus Mass Spectrometer (Thermo Scientific). Samples (10μl) were loaded onto a C18 trap column and washed for 5 minutes with 0.1% formic acid. Peptides were resolved using a gradient (170 min, 0.3μl/min) of buffer A (0.1% formic acid) and buffer B (80% acetonitrile in 0.08% formic acid): 5% buffer B for 5 min, 5-35% buffer B for 125 min, 35-98% buffer B for 2 min, 98% buffer B for 20 min, 98-2% buffer B for 1 min and 2% buffer B for 17 min. Peptides, initially trapped on an Acclaim PepMap 100 C18 colum (100μm x 2 cm, Thermo Scientific), were separated on an Easy-Spray PepMap RSLC C18 column (75μm x 50 cm, Thermo Scientific), and finally transferred to a Q Exactive Plus Mass Spectrometer via an Easy-Spray source with temperature set at 50°C and a source voltage of 2.0kV. For the identification of peptides, a top 15 method (1 MS plus 15 MS^2^, 150 min acquisition) consisting of full scans and mass range (m/z) between 350 to 1600 (m/z) for MS search and 200 to 2000 (m/z) for MS^2^ search was used. For the MS scan the Q Exactive Plus Mass Spectrometer was operated in a data dependent acquisition mode, resolution of 70,000 with a lock mass set at 445.120024 and max fill time of 20 ms. For the MS^2^ scan Q Exactive Plus Mass Spectrometer was operated in a centroid mode, resolution of 15,000 with isolation window = 1.4 (m/z), normalized collision energy = 27, max fill time of 100 ms and dynamic exclusion of 45.0 sec.

### Mass spectrometry data analysis

Q Exactive Plus Mass Spectrometer .RAW files were analyzed, and peptides and proteins quantified using MaxQuant (Cox and Mann, 2008), using the built-in search engine Andromeda (Cox et al., 2011). All settings were set as default, except for the minimal peptide length of 5, and Andromeda search engine was configured for the UniProt *Homo sapiens* protein database (release date: 2018_09). Peptide and protein ratios only quantified in at least two out of the three replicates were considered, and the p-values were determined by Student’s t test and corrected for multiple testing using the Benjamini–Hochberg procedure (Benjamini and Hochberg, 1995).

### GO Analysis

DAVID GO analysis tool (Huang da et al., 2009; Huang et al., 2007) was used to find statistically over-represented gene ontology (GO) categories.

### Inhibition of Ser727 STAT3 phosphorylation

Resting CD4^+^ T cells were labeled with anti-CD4-FiTC antibody (Biolegend, #357406) and isolated from human PBMCs by magnetic activated cell sorting (MACS, Miltenyi) using anti-FiTC microbeads (Miltenyi, Cat#130-048-701) following manufacturer instructions. Then, CD4^+^ T cells were activated for three days with ImmunoCult Human CD3/CD28 T Cell Activator (StemCell, #10971) following manufacturer instructions in the presence of 20 ng/mL IL2 (Novartis, #709421). After activation, cells were expanded for 5 days in the presence of 20ng/mL IL2. Then, cells were starved of IL2 for 24 hours before stimulation with 10nM HyIL-6 in the presence or absence of different inhibitors [2μM Tofacitinib (Stratech, #S2789-SEL), 2μM Rapamycin (Stratech, #S1039-SEL), 2μM Torin1 (Tocris, #4247), 2μM CHIR-99021 (Stratech, #G09-901B-SGC), 2μM PD184352 (Stratech, #S1020-SEL), 2μM Roscovitine (Calbiochem, #557360), 2μM GDC0941 (abCam, #ab141352), 2μM PP2 (Stratech, #S7008-SEL), 2μM VX745 (Tocris, #3915), 2μM JAK inhibitor VII (Calbiochem, #796041-65-1), 2μM AX15836 (Tocris, #5843), 2μM AZD8055 (Stratech, #A8214-APE), 2μM KU0063794 (Tocris, #3725), 2μM KU55933 (Stratech, #A4605-APE), 2μM KU57788 (Stratech, #A8315-APE), 2μM CBI-2347 (a kind gift of Boehringer Ingelheim), 2uM MSC2530818 (Stratech, #S8387-SEL), 2μM CDK9 inhibitor II (Calbiochem, #140651-18-9), 2μM NVP2 (Tocris, #6535), 2μM THZ531 (Stratech, #A8736-APE), 2μM MFH-2-90-1 (a kind gift of Dr. Greg Findlay, University of Dundee, UK) and 2μM Flavopiridol (Stratech, #S2679-SEL)] as indicated.

### Western blotting

Cells were rinsed in ice-cold PBS then lyzed in RIPA buffer (Thermo Scientific) supplemented with protease inhibitor cocktail (ROCHE), 5 mM sodium fluoride, 2 mM sodium orthovanadate and 0.2 mM PMSF incubating on ice for 15 min. Lysates were cleared by centrifugation at 20,000 g for 15 min at 4°C then protein concentrations determined using Coomassie Protein Assay Kit (Thermo Scientific, UK). For each sample, 30 μg of total protein were separated on 4-12% Bis-Tris polyacrylamide gels (NuPAGE, Invitroge) in MES SDS running buffer then blotted onto Protran 0.2 mM Nitrocellulose (GE Healthcare, UK). Membranes were probed with 1:1000 dilution of the appropriate primary antibody anti-pSer727-STAT3 (Cell Signaling, #9134S) anti-p-Tyr705-STAT3 (Cell Signaling, #9145S), anti-total-STAT3 (Cell Signaling, #9139S), anti-total-STAT1 (Transduction Laboratories, #G16920), anti-Lamin A (Santa Cruz, #sc-20680), anti-GAPDH (Cell Signaling, #2118S), anti-pS933-NFκβ (Cell Signaling, #4806S), anti-pS473-AKT (Cell Signaling, #4060S), anti-pT389-S6K (Cell Signaling, #9234S), anti-pS177-IKKαβ (Cell Signaling, #2078S), anti-pS376-MSK1 (Cell Signaling, #9591S), anti-pPan(T514)-PKC (Cell Signaling, #9379S), anti-pY783-PLCγ1 (Cell Signaling, #2821S), anti-pS21/S9-GSK3α/β (Cell Signaling, #9331S), anti-pT202/Y204-ERK1/2 (Cell Signaling, #4370L), anti-pT180/Y182-P38 (Cell Signaling, #9215S), anti-pT183/Y185-JNK (Cell Signaling, #9251S). 1:5000 dilution of donkey anti-rabbit-HRP (Stratech, 711-035-152-JIR) or donkey anti-mouse-HRP (Stratech, 715-035-150-JIR) as the secondary antibody. Immobilon Western Chemiluminescent HRP substrate (Millipore, UK) was used for visualization.

### Subcellular fractionation

Resting CD4^+^ T cells were labeled with anti-CD4-FiTC antibody (Biolegend, #357406) and isolated from human PBMCs by magnetic activated cell sorting (MACS, Miltenyi) using anti-FiTC microbeads (Miltenyi, Cat#130-048-701) following manufacturer instructions. Then, CD4^+^ T cells were activated for three days with ImmunoCult Human CD3/CD28 T Cell Activator (StemCell, #10971) following manufacturer instructions in the presence of 20 ng/mL IL2 (Novartis, #709421). After activation, cells were expanded for 5 days in the presence of 20ng/mL IL2. Then, cells were starved of IL2 for 24 hours before stimulation as indicated. Cells were subsequently washed three times with ice-cold PBS and subcellular fractionation was carried out using a Cell Fractionation kit following manufacturer instructions (Cell Signaling, #9038).

### Proximity ligation assay

Resting CD4^+^ T cells were labeled with anti-CD4-FiTC antibody (Biolegend, #357406) and isolated from human PBMCs by magnetic activated cell sorting (MACS, Miltenyi) using anti-FiTC microbeads (Miltenyi, Cat#130-048-701) following manufacturer instructions. Then, CD4^+^ T cells were activated for three days with ImmunoCult Human CD3/CD28 T Cell Activator (StemCell, #10971) following manufacturer instructions in the presence of 20 ng/mL IL2 (Novartis, #709421). After activation, cells were expanded for 5 days in the presence of 20ng/mL IL2. Then, cells were starved of IL2 for 24 hours before stimulation as indicated and 10^5^ cells were used per experiment. Cells were attached to coverslips by incubating them at 37°C for 1 hour in PBS, then PBS was replaced with RPMI supplemented with 10% FBS and cells stimulated as described. After stimulation, cells were fixed with 2% formaldehyde for 10 minutes at RT, permeabilized with ice-cold methanol for 20 minutes on ice and stained with anti-STAT3 (Cell Signaling, #9139S) and anti-CDK8 (Invitrogen, #PA1-21780) or anti-STAT3 (Cell Signaling, #9139S) and anti-CDK9 (Cell Signaling, #2316S) for Proximity Ligation Assays following manufacturer instructions (Sigma, #DUO92008).

### Chromatin immunoprecipitation by sequencing (ChIP-Seq)

In vitro polarized human Th-1 cells were expanded in the presence of IL-2 for 10 days and cells were then washed with complete media and rested for 24 hr starvation in the absence of IL-2, these cells were then either not-stimulated (control) or stimulated with IL-6 or different IL-6 variants for 1 hr, cells were then immediately fixed with 1% methanol-free formaldehyde (Formaldehyde 16%, Methanol-Free, Fisher Scientific, PA, USA) at room temperature for 10mn with gentle rocking cells were then washed twice with cold PBS. For each STAT3 ChIP-seq library sample, approximately 10 × 106 cells were used and the fixed cell palettes were kept at −80°C prior to further processing. The ChIPseq experiments were performed as previously described (Liao et al., 2008) with some modification as described below. In brief, the frozen cell pellets were thawed on ice and washed once with 1 mL cold PBS by centrifugation at 5000 RPM for 5 min, the resulting cell pellets were re-suspended in 500 uL of lysis buffer (1X PBS, 0,5% Triton X-100, cOmplete EDTA-free protease inhibitor cocktail, Roche Diagnostics, Basel, Switzerland) and incubated for 10 min on ice, followed by a 5 min centrifugation at 5000 RPM. Then the pellets were washed once with 1 mL of sonication buffer (1X TE, 1: 100 protease inhibitor cocktail), re-suspended in 750 uL of sonication buffer (1X TE, 1: 100 protease inhibitor cocktail and 0,5 mM PMSF) and sonicated for 20 cycles (on-20sec and off-45sec) on ice using VCX-750 Vibra Cell Ultra Sonic Processor (Sonics, USA). The sonicated lysates were centrifuged 20 min at 14000 RPM and the clear lysate supernatants were collected and incubated with 30 uL of Protein-A Dynabeads (ThermoFisher, USA) that were pre-incubated with incubated with 10 ug of anti-STAT3 antibody (anti-Stat3, 12640S, Cell Signaling Technology) at 4°C overnight with gentle rotation. Next day, the beads were washed 2 times with RIPA-140 buffer (0.1% SDS, 1% Triton X-100, 1 mM EDTA, 10 mM Tris pH 8.0, 300 mM NaCl, 0.1% NaDOC), 2 times with RIPA-300 buffer (0.1% SDS, 1% Triton X-100, 1 mM EDTA, 10 mM Tris, 300 mM NaCl, 0.1% NaDOC), 2 times with LiCl buffer (0.25 mM LiCl, 0.5% NP-40, 1 mM EDTA, 10 mM Tris pH 8.0, 0.5% NaDOC), once with TE-0,2% Triton X-100 and once with TE buffer. Crosslinks were reversed by incubating the bound complexes in 60 uL TE containing 4.5 uL of 10% SDS and 7.5 uL of 20 mg/mL of proteinase K (Thermofisher, USA) at 65°C overnight for input samples, we used 6 uL of 10% SDS and 10 μL of 20 mg/mL of proteinase K. Then, the supernatants were collected using a magnet and beads were further washed one in TE 0.5M NaCl buffer. Both supernatants were combined, and DNA was extracted with phenol/chloroform, followed by precipitation with ethanol and re-suspended in TE buffer. The library was constructed following the manufacturer protocol of the KAPA LTP Library Preparation Kit (KAPA Biosystems, Roche, Switzerland). ChIP DNA libraries were ligated with the Bioo scientific barcoded adaptors (BIOO Scientific, Perkin Elmer, USA) with T4 DNA ligase according to KAPA LTP library preparation protocol and the ligated ChIP DNA libraries were purified with 1.8x vol. Agencourt AMPure XP beads and PCR amplified using KAPA hot start High-Fidelity 2X PCR Master Mix and NextFlex index primers (Bioo Scientific, PerkinElmer) for 12 cycle by following thermocycler cycles: 30 s hot start at at 98°C, followed by 12 cycle amplification [98°C for 10 s, 60°C for 30 s and 72°C for 30 s] and final extension at 72°C for 1 min. The amplification and quality of the ChIPseq libraries were checked by running 10% of the samples in E-Gel Agarose Gels with SYBR Safe DNA Gel Stain (ThermoFisher Scientific, USA), and if necessary, samples were reamplified additional four cycles using the same thermocycler protocol described above. Then, the libraries were purified and size-selected using Agencourt AMPure XP beads (1.25x vol. to remove short fragments. The concentration of ChIP-DNA libraries was measured by Qubit-4 fluorometer (ThermoFisher, USA) and equal amounts of each sample were pooled and 50 bp paired-end reads were sequenced on an Illumina 4000 platform by GENEWIZ technology (GENEWIZ, USA).

### RNA-sequencing

For RNA-seq library preparation, in vitro polarized human Th-1 cells either not stimulated or stimulated with HyIL-6 in the presence or absence of 2μM MSC2530818 variants at 37°C for 6 hr, total RNA was extracted and RNAseq libraries were prepared by Edinburg Sequencing Core facility.

### ChIP-Seq data analysis

The quality of libraries was inspected using FastQC v0.11.8. All sequencing reads were aligned to human reference genome (GRCh37; hg19) using bowtie v1.2.2 (Langmead et al., 2009) with default pair-end alignment settings and additional parameters ‘--chunkmbs 1000 S -m 1’. The index for reference genome was constructed by using ‘bowtie-build’ with default parameters. Sorting and indexing of the aligned reads were conducted by Samtools v1.9 (Li et al., 2009). The genome-wide binding profiles (i.e. bigWig files) were generated by bamCoverage v3.2.0 (Ramirez et al., 2016) using parameters ‘--normalizeUsing BPM -- minMappingQuality 30 --ignoreDuplicates --extendReads 250 --blackListFileName hg19.blacklist.bed’. The binding profiles were visualized using IGV genome browser v2.7.0 (Robinson et al., 2011). Binding peaks were called by ‘callpeaks’ procedure from MACS2 v2.1.2 (Zhang et al., 2008) using default parameters except ‘-f BAMPE --nomodel -t treatment -c input’. The identified peaks were further screened against ‘hg19 blacklisted’ genomic regions, mitochondrial DNA, and pseudo-chromosomes. *De novo* motif findings were performed in 200 bp surrounding the summit of n = 500 top bound regions using MEME Suite v5.0.2 (Bailey et al., 2009) with default parameters except ‘-maxsize 10000000 -dna -mod zoops -nmotifs 10’. *De novo* motifs were compared against all JASPAR known motifs by TOMTOM (Gupta et al., 2007). STAT3 shared bound regions in HyIL6 (n=540) or HyIL-6+MSC (n=2585) stimulated cells were generated by the intersection between bound regions from n=3 donor using BEDTools (Quinlan and Hall, 2010). STAT3 binding intensity in shared bound regions was calculated by UCSC bigWigAverageOverBed v2 with default parameters and the mean signal intensity was visualized by PRISM v8.4.0. The shared STAT3 bound regions were annotated with the nearest gene by ‘annotatePeaks’ from HOMER v4.10 (Gupta et al., 2007), yielding 475 unique genes. Statistical analyses were performed using the Two-tailed parametric and non-parametric tests as appropriate.

### RNA-Seq data analysis

The quality of libraries was inspected by FastQC v0.11.8. The expression level of mRNA in each library was quantified by ‘rsem-calculate-expression’ in RSEM v1.3.1 (Li and Dewey, 2011) using default parameters except ‘--bowtie-n 1 --bowtie-m 100 -seed-length 28 --paired-end’. The bowtie index required by RSEM software was generated by ‘rsem-prepare-reference’ on all RefSeq genes, downloaded from UCSC table browser on April 2017. EdgeR v3.24.0 (Robinson et al., 2010) package was used to normalize gene expression among all libraries and identify differentially expressed genes among samples with following constraints: fold change ≥ 1.5, p-value≤0.05. Scatter and bar plots were drawn by Datagraph v4.5 and PRISM v8.4.0, respectively. Geneset enrichment analysis was performed by GSEA v4.0.3 (Subramanian et al., 2005) with default parameters except ‘-collapse No_Collapse -permute gene_set’. Pathway analysis of differentially expressed genes was performed by Metascape (Zhou et al., 2019) on all GO terms related to biological processes, KEGG Pathways, BioCarta Gene Sets and Hallmark Gene Sets.

### T cells population differentiation

Resting CD4^+^ T cells isolated as described above were activated under Th-1, Th-17 or Tregs polarizing conditions. Briefly, resting human CD4^+^ T cells freshly isolated were activated using ImmunoCult Human CD3/CD28 T Cell Activator (StemCell, Cat#10971) following manufacturer instructions for 3 days in the presence of the cytokines required for the different CD4^+^ T cells populations: Th-1 (IL-2 (20 ng/ml), anti-IL-4 (10 ng/ml), IL-12 (20 ng/ml)), Th-17 (IL-1β (10 ng/ml, R and D Systems, Cat#201-LB/CF), IL-23 (10 ng/ml, R and D Systems, Cat#1290-IL), anti-IL-4 (10 ng/ml), anti-IFNγ (10 ng/ml, BD Biosciences, Cat#554698)) or Tregs (IL2 (20 ng/ml), TGF-β (5 ng/ml, Peprotech, Cat#100–21), anti-IL-4 (10 ng/ml), anti-IFNγ (10 ng/ml)) in the presence or absence of saturating concentrations of the different variants of IL-6 described in this manuscript. After three days of priming, cells were expanded for another 5 days in the presence of IL-2 (20 ng/ml). Th-1 and Th-17 cells were restimulated for 6 hr in the presence of PMA (100 ng/ml, Sigma, Cat#P8139), Ionomycin (1 μM, Sigma, I0634) and Brefeldin A (5 μg/ml, Sigma, B7651) before FACS analysis. In all cases cells were fixed with 2% formaldehyde and prepared to be analysed by FACS. Cells were then permeabilised with Saponin 2% in PBS for 20 min at room temperature and then stained in Saponin 2% in PBS with the appropriate antibodies: Th-1 (anti-CD3-BV510 (1:100, Biolegend, Cat#300448), anti-CD4-PE (1:100, Biolegend, Cat#357404), anti-CD8-AF700 (1:100, Biolegend, Cat#300920), anti-IFNγ (1:100, Biolegend, Cat#502217)), Th-17 (anti-CD3-BV510, anti-CD4-PE, anti-CD8-AF700, anti-IL17A-APC (1:100, Biolegend, Cat#512334)) and Tregs (anti-CD3-BV510, anti-CD4-PE, anti-CD8-AF700, anti-CD25-APC (1:100, Biolegend, Cat#302610), anti-FoxP3-AF488 (1:100, Biolegend, 320012)) and analysed in a CytoFLEX S (Beckman Coulter).

## SUPPLEMENTARY FIGURE LEGENDS

**Supplementary Figure 1.**
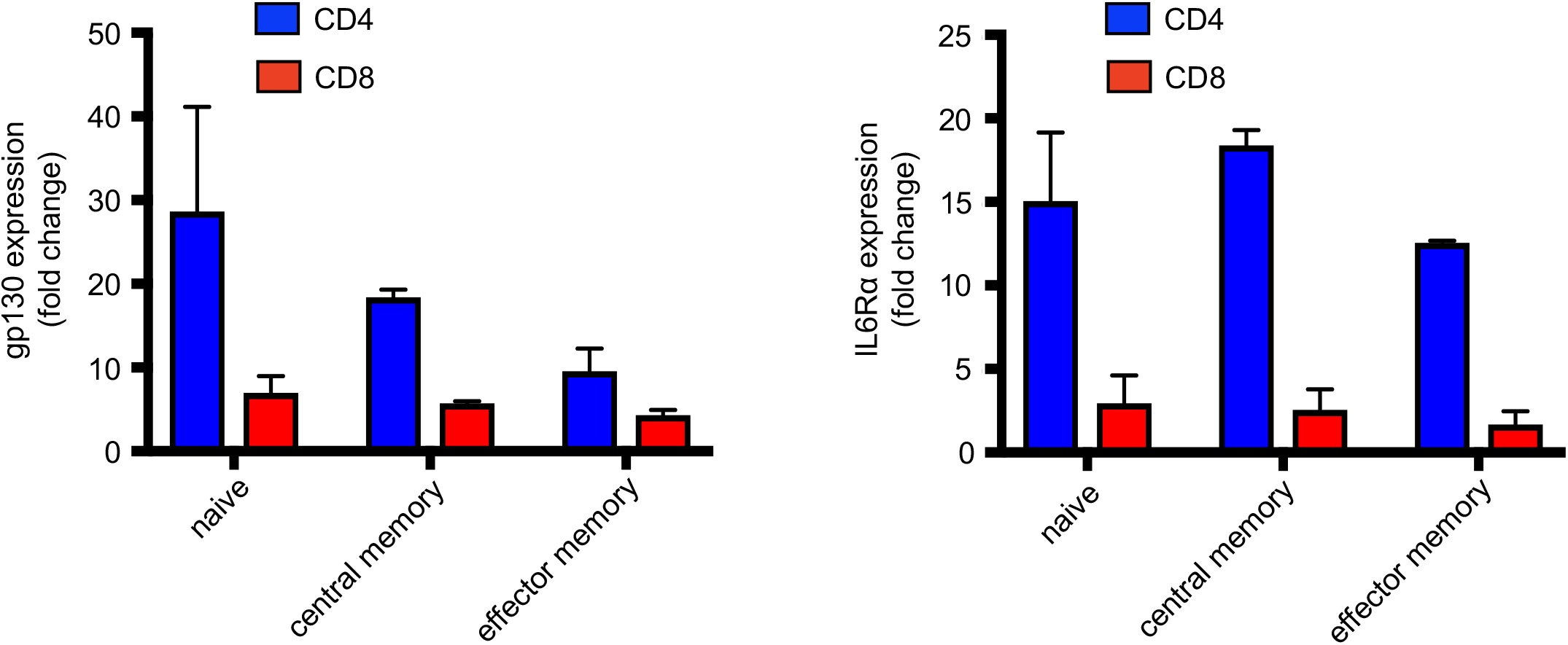
Expression levels of gp130 and IL-6Rα in different subpopulations of human T cells. Levels of expression of gp130 (left panel) and IL-6Rα (right panel) expressed as fold change in different population of resting primary human CD4^+^ and CD8^+^ T cells.

**Supplementary Figure 2.**
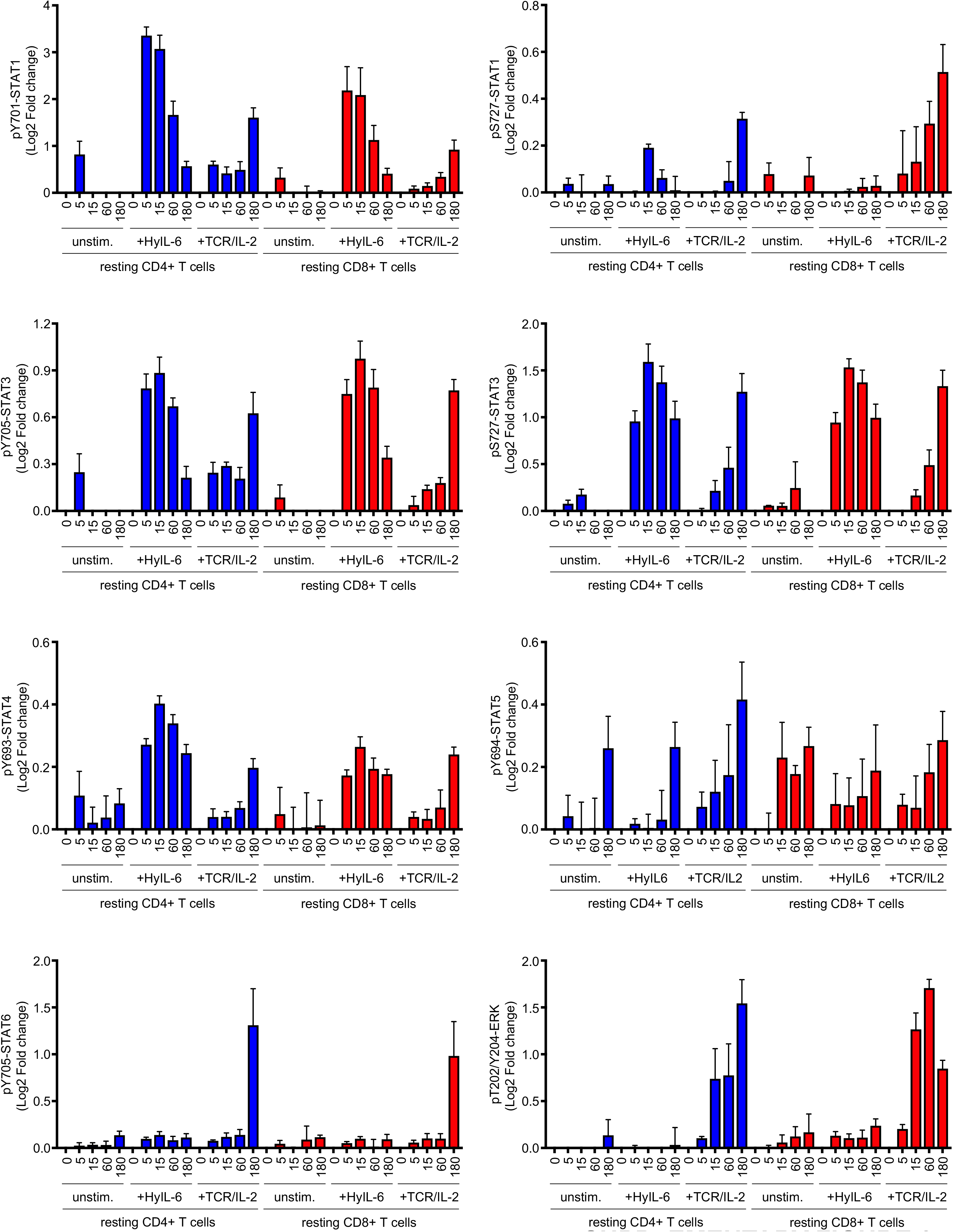
Phospho-FLOW analyses of the HyIL-6 signalosome in resting human primary CD4^+^ and CD8^+^ T cells: Time-course. Time-course data of the changes induced by HyIL-6 in the phosphorylation state of the main signaling pathways in resting human primary CD4^+^ and CD8^+^ T cells unstimulated, treated with HyIL-6 or with anti-CD3+Interleukin-2, as shown in Figure 1C.

**Supplementary Figure 3.**
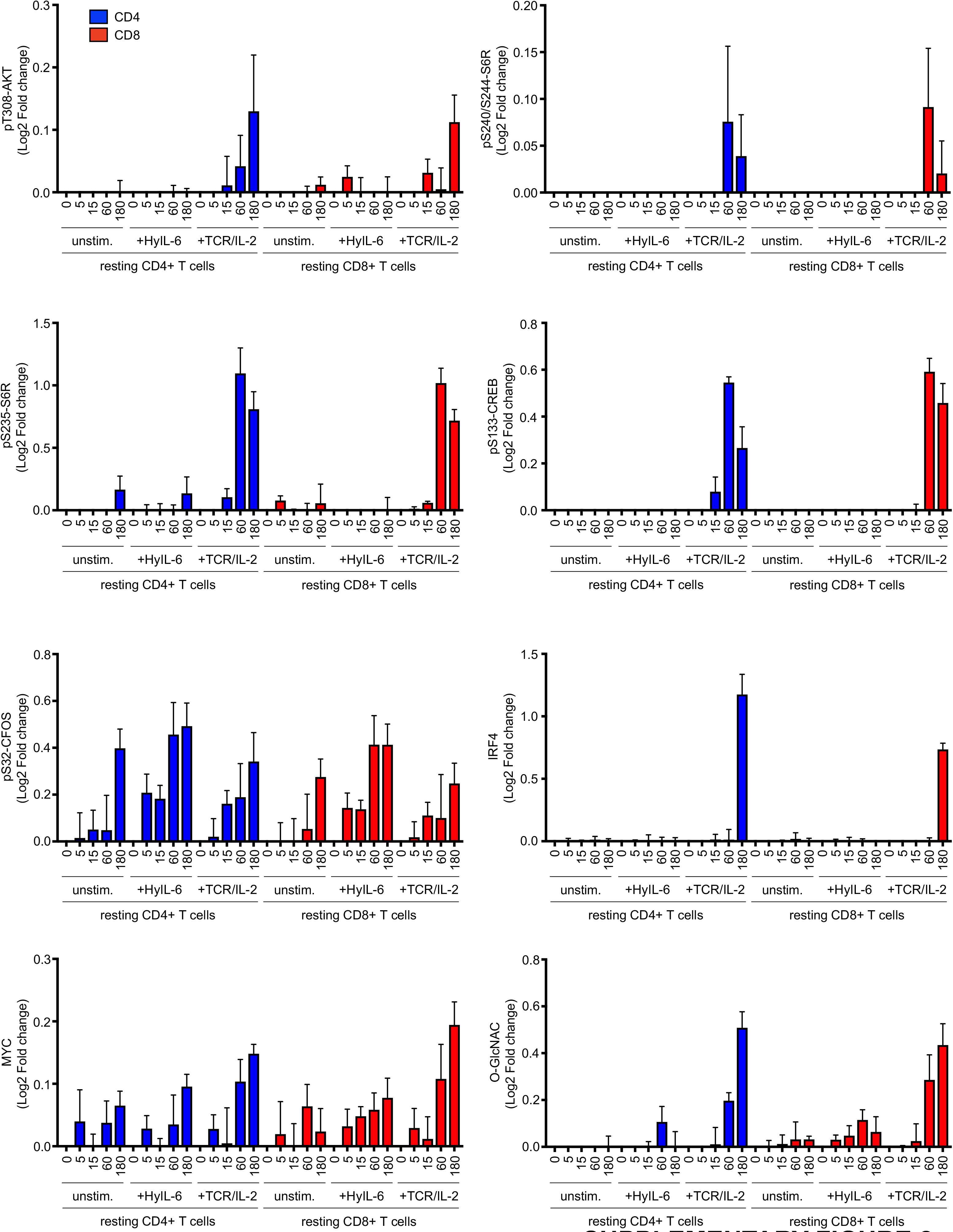
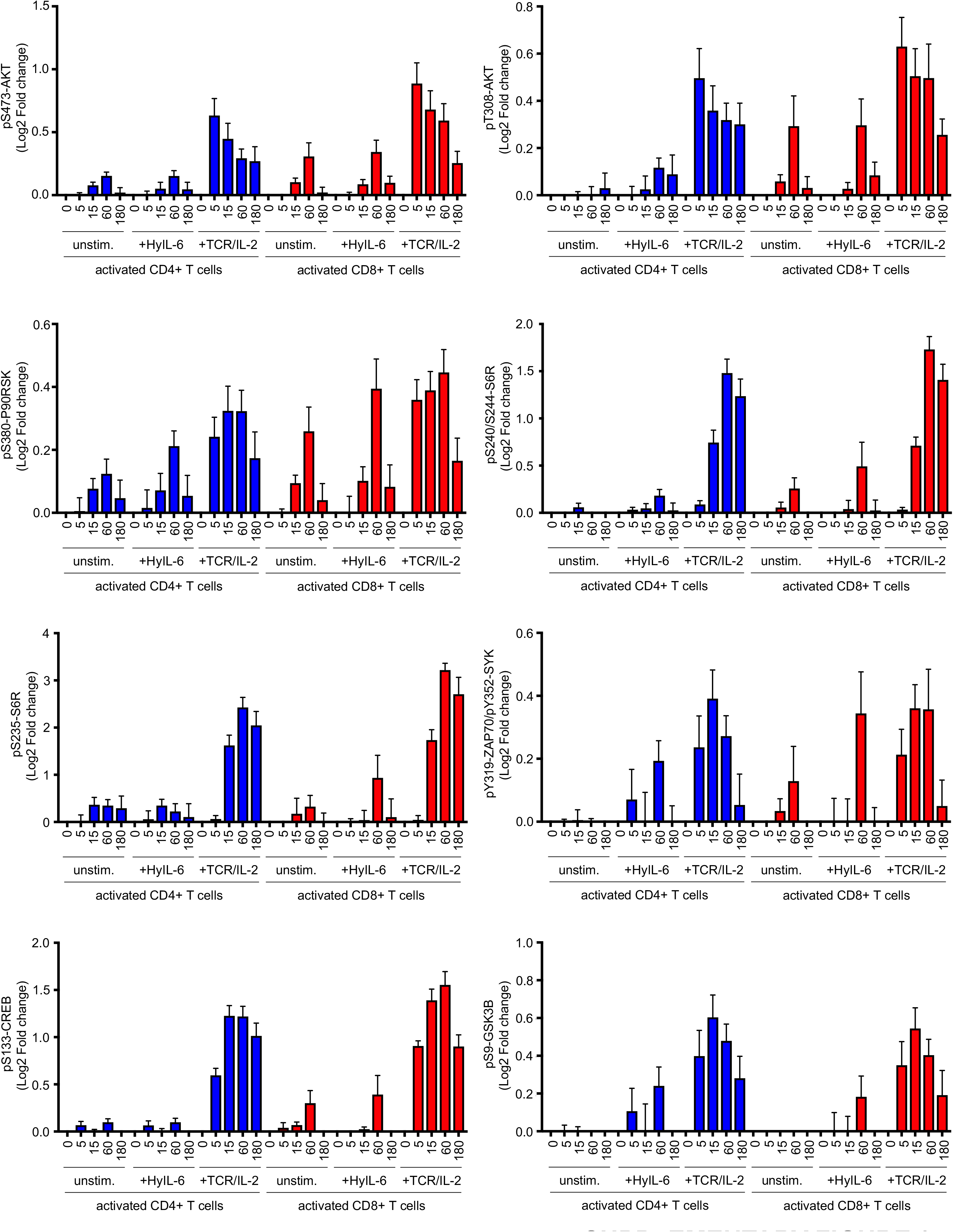
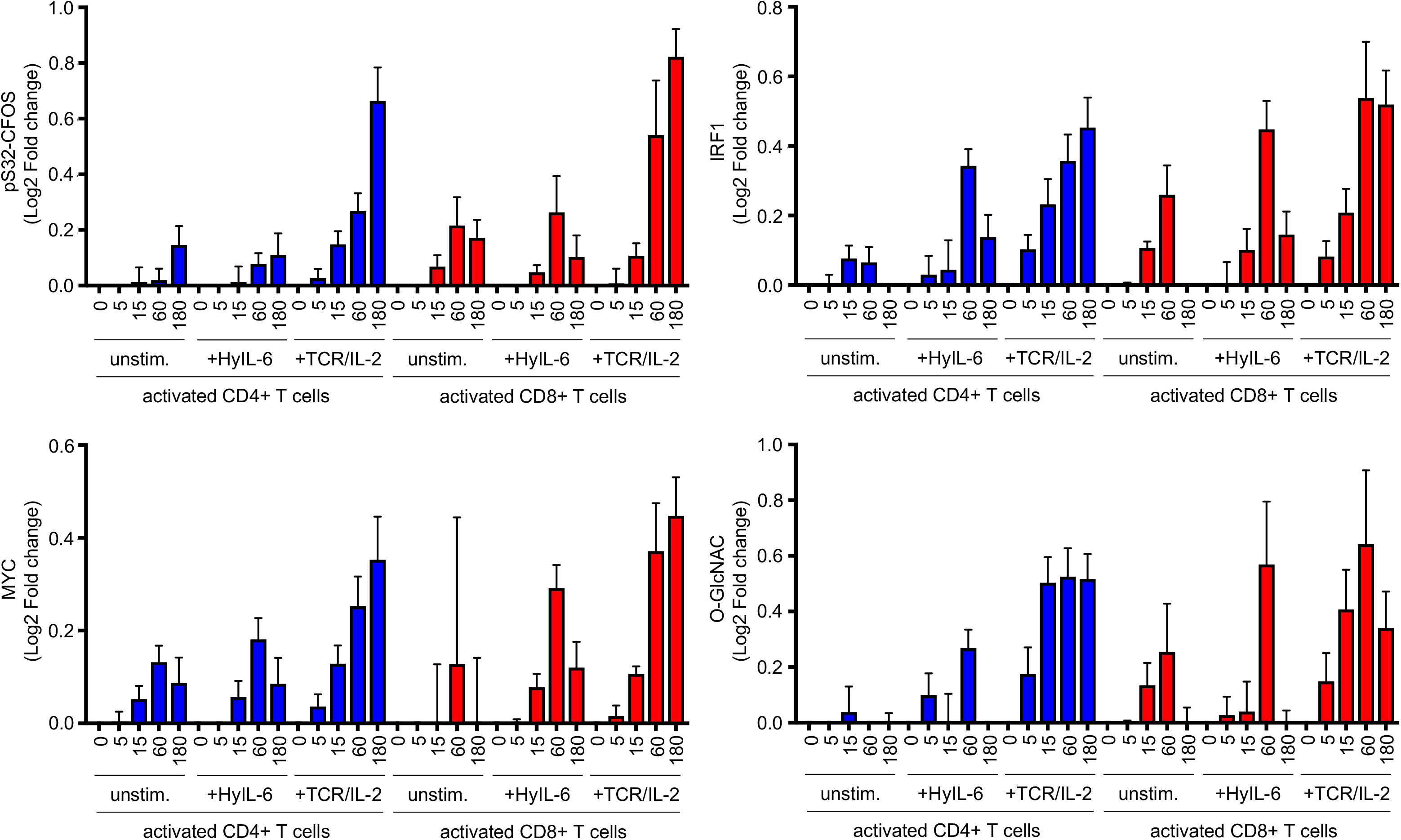
Phospho-FLOW analyses of the HyIL-6 signalosome in activated human primary CD4^+^ and CD8^+^ T cells: Time-course. Time-course data of the changes induced by HyIL-6 in the phosphorylation state of the main signaling pathways in activated human primary CD4^+^ and CD8^+^ T cells unstimulated, treated with HyIL-6 or with anti-CD3+Interleukin-2, as shown in Figure 1D.

**Supplementary Figure 4.**
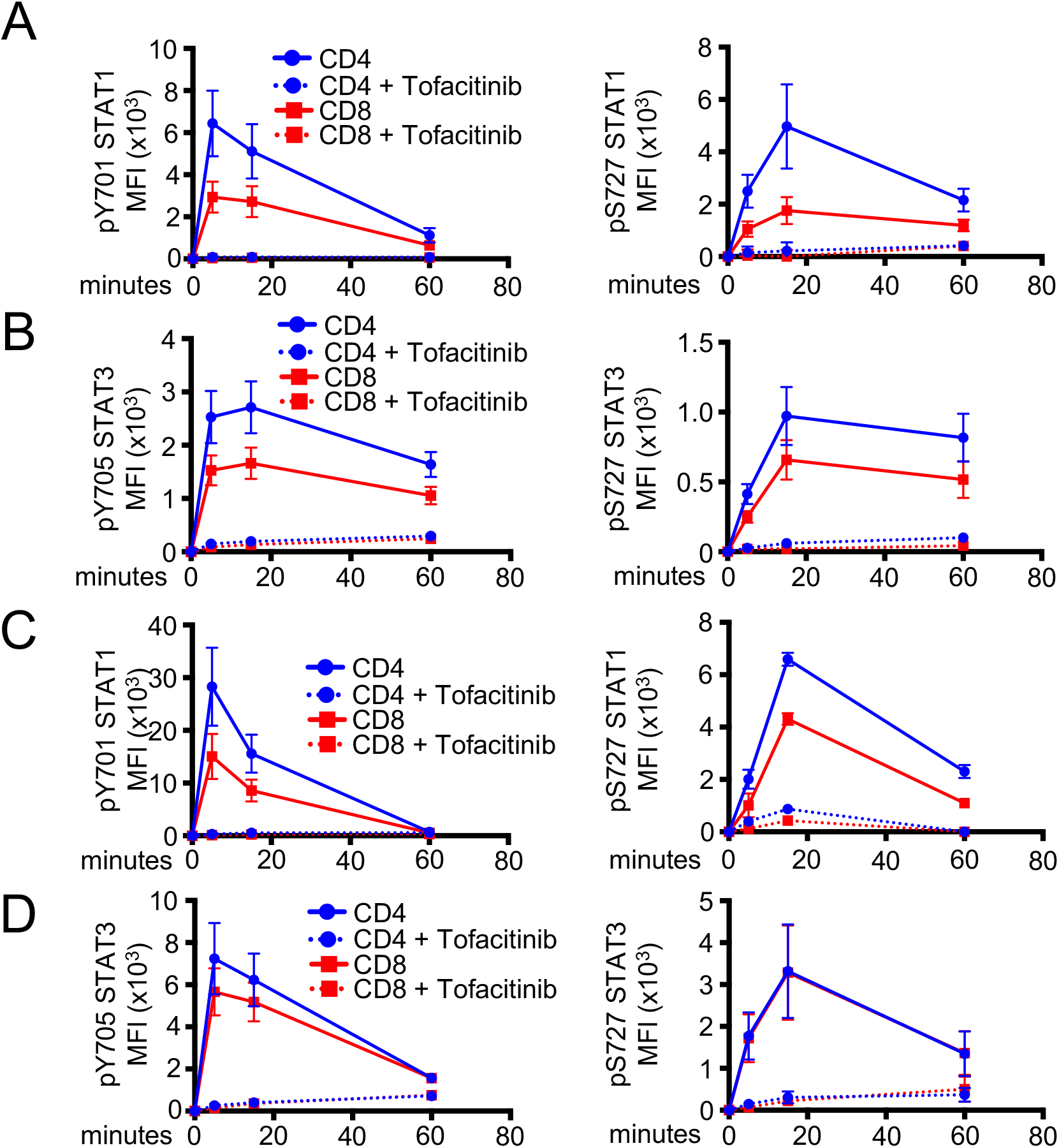
Effect of Tofacitinib in the HyIL-6-induction of Tyr and Ser phosphorylation in STAT3 and STAT1 in resting and activated primary human CD4^+^ and CD8^+^ T cells. **A)** Immunoblotting of some of the main pathways regulated by Interlekin-6 as described in the literature in resting primary human CD4^+^ T cells not stimulated, treated with HyIL-6 or anti-CD3+Interleukin-2. **B)** Time-course of STAT1 Tyr701 (left panel) and Ser727 (right panel) phosphorylation induced by Interleukin-6 stimulation in the presence (dash line) or absence (solid line) of 2μM Tofacitinib in resting primary human CD4^+^ and CD8^+^ T cells. **C)** Time-course of STAT3 Tyr705 (left panel) and Ser727 (right panel) phosphorylation induced by Interleukin-6 stimulation in the presence (dash line) or absence (solid line) of 2μM Tofacitinib in resting primary human CD4^+^ and CD8^+^ T cells. **D)** Time-course of STAT1 Tyr701 (left panel) and Ser727 (right panel) phosphorylation induced by Interleukin-6 stimulation in the presence (dash line) or absence (solid line) of 2μM Tofacitinib in activated primary human CD4^+^ and CD8^+^ T cells. **E)** Time-course of STAT3 Tyr705 (left panel) and Ser727 (right panel) phosphorylation induced by Interleukin-6 stimulation in the presence (dash line) or absence (solid line) of 2μM Tofacitinib in activated primary human CD4^+^ and CD8^+^ T cells.

**Supplementary Figure 5.**
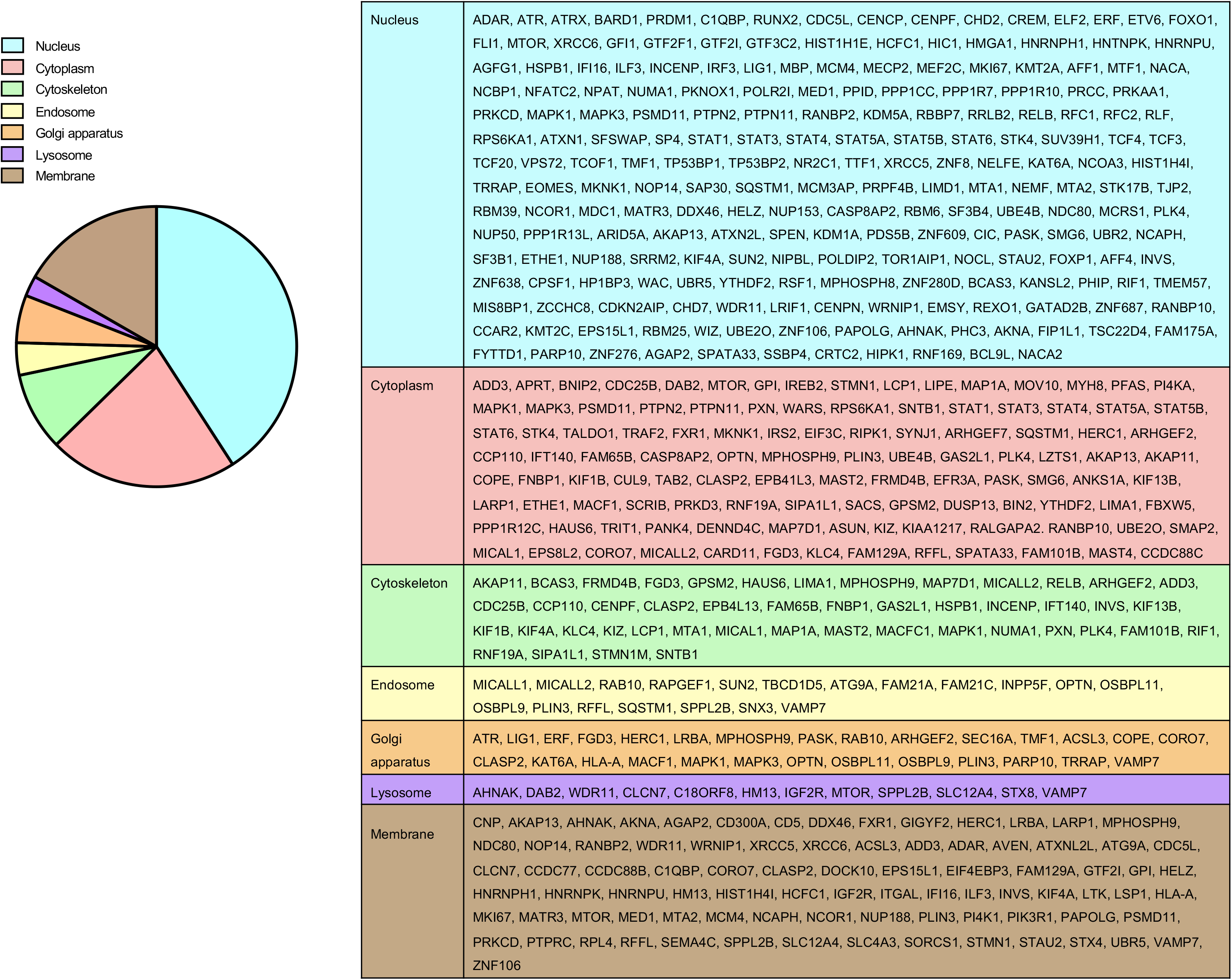
Cellular location of the proteins regulated by HyIL-6 in primary human Th-1 cells. **A)** Gene Ontology analysis of the cellular location of the proteins regulated by phosphorylation in human primary CD4^+^ Th-1 cells in response to HyIL-6.

**Supplementary Figure 6.**
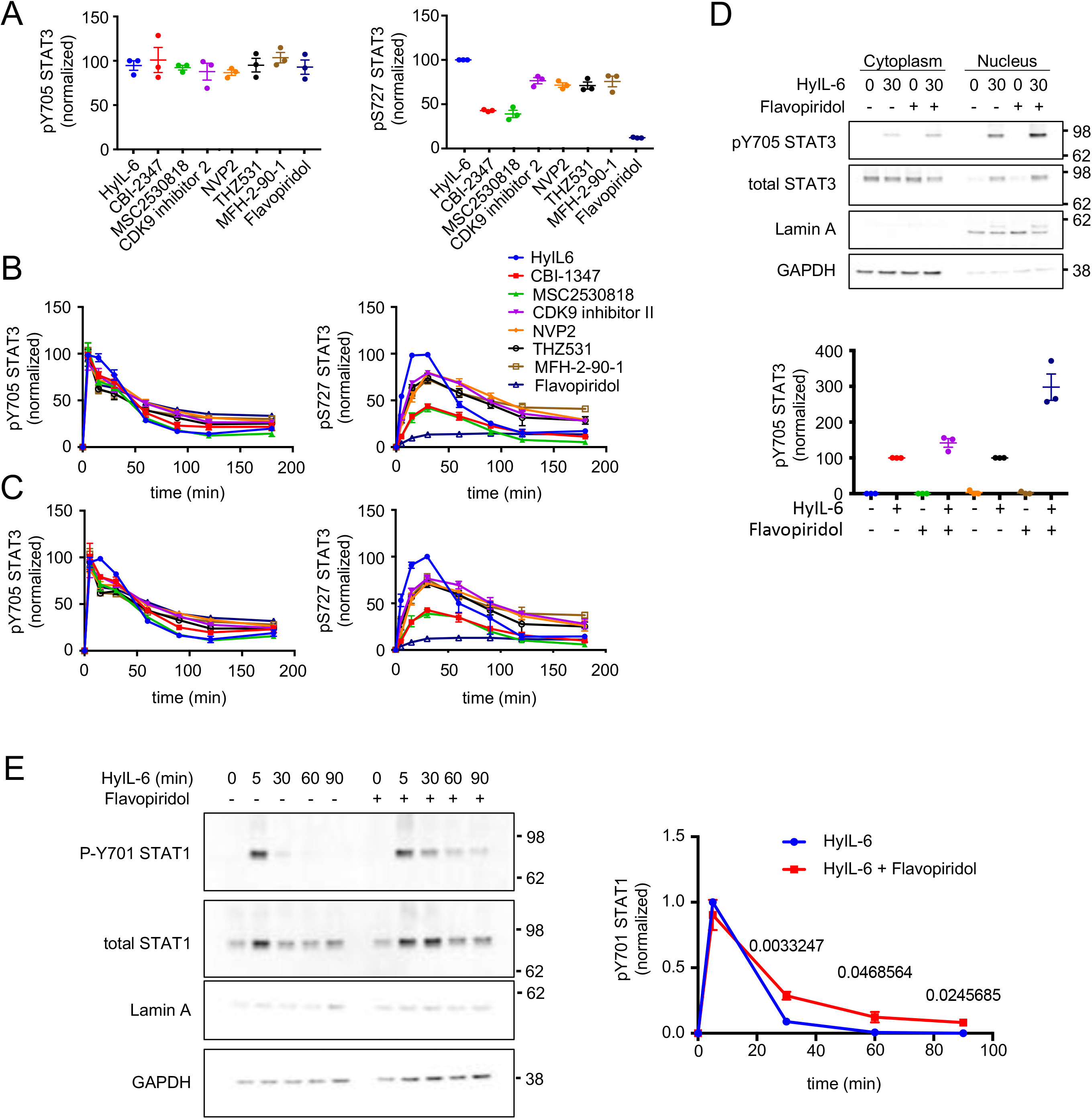
Role of CDKs in the regulation of Ser727 STAT3 and STAT1 phosphorylation. **A)** Effect of different CDK inhibitors on the STAT3 Tyr705 (left panel) and STAT3 Ser727 (right panel) phosphorylation induced by HyIL-6 in human primary CD8^+^ T cells. **B)** Kinetics of the HyIL-6-induced STAT3 Tyr705 (left panel) and Ser727 (right panel) phosphorylation in the presence of different CDK inhibitors in human primary CD4^+^ T cells. **C)** Kinetics of the HyIL-6-induced STAT3 Tyr705 (left panel) and Ser727 (right panel) phosphorylation in the presence of different CDK inhibitors in human primary CD8^+^ T cells. **D)** Immunoblotting analysis of the level of phosphorylation of STAT3 Tyr705 in the cytoplasmic and nuclear fraction of human primary CD4 T cells after 30 minutes stimulation with HyIL-6 in the absence or presence of Flavopiridol. Quantitative analysis showing the normalized data of three independent biological samples is shown alongside. **E)** Time-course of STAT1 Tyr701 phosphorylation in nuclear fraction of human primary CD4^+^ cells stimulated with HyIL-6 in the absence (blue line) or presence (red line) of 2μM Flavopiridol. Quantitative analysis showing the normalized data of three independent biological samples is shown alongside. **F)** Timecourse of RPB1 Ser2 and Ser5 phosphorylation upon HyIL-6 stimulation in activated primary human Th-1 cells in the absence or presence of MSC2530818. Quantitative analysis showing the normalized data of three independent biological samples is shown alongside.

**Supplementary Figure 7.**
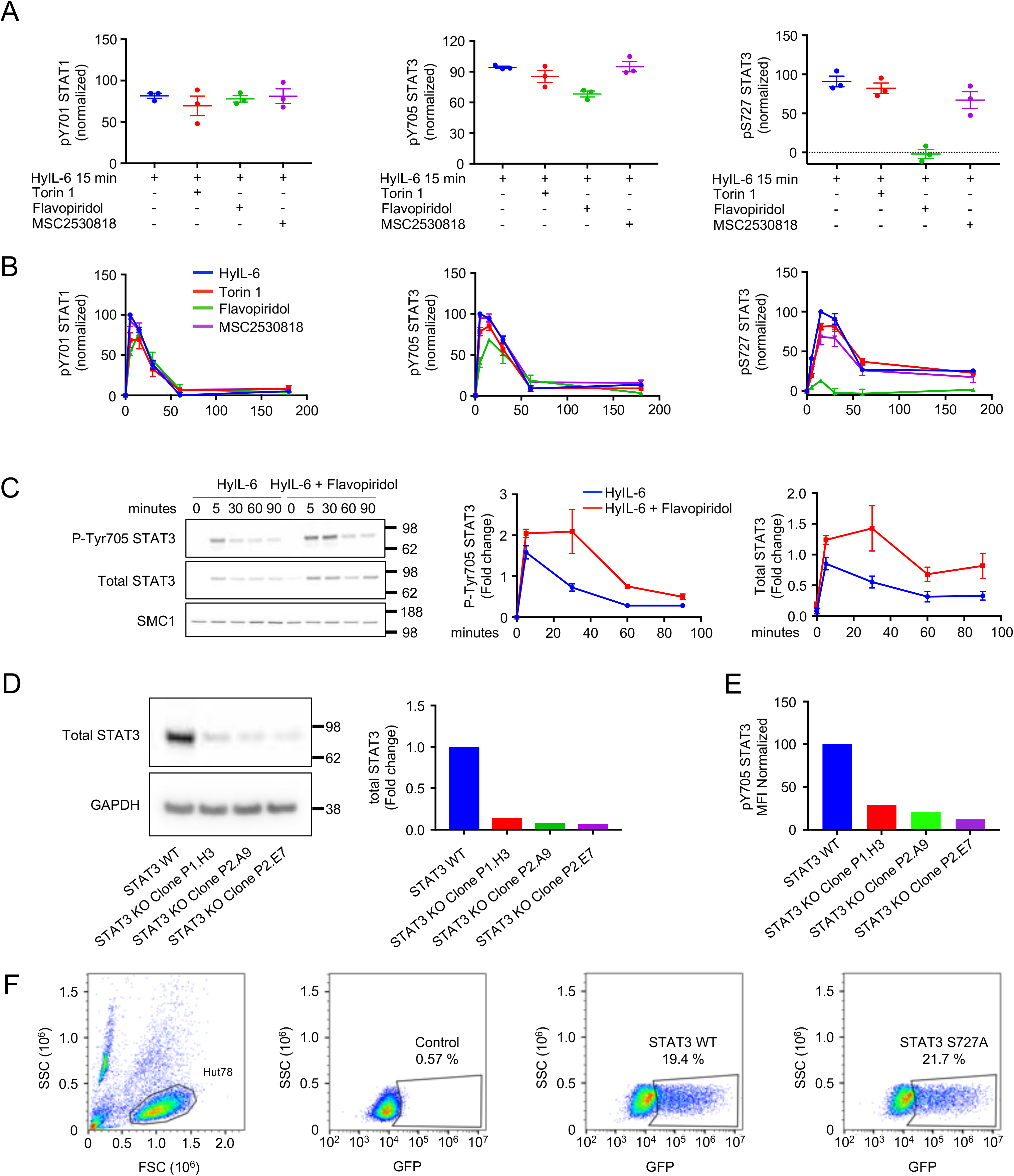
Regulation of Ser727 STAT3 phosphorylation in Hut78 cells. **A)** Effect of Torin1, Flavopiridol or MSC2531818 on the STAT1 Tyr701 (upper panel), STAT3 Tyr705 (middle panel) and STAT3 Ser727 (lower panel) phosphorylation induced by HyIL-6 in human primary CD4^+^ T cells. **B)** Time-course of STAT1 Tyr701 (upper panel), STAT3 Tyr705 (middle panel) and STAT3 Ser727 (lower panel) phosphorylation induced by HyIL-6 in human primary CD4^+^ T cells in the presence or absence of Torin1, Flavopiridol or MSC2530818. **C)** STAT3 immunoblot of Hut78 WT cells vs Crispr/CAS9 generated STAT3 knock-downs (KDs) in Hut78. Quantitation of the levels of STAT3 are shown in the graph. **D)** FACS analysis of the level of STAT3 Tyr705 phosphorylation in Hut78 WT and the different STAT3 KDs Hut78 cell lines upon 15 min HyIL-6 stimulation. **E)** FACS analysis of the expression of STAT3 WT-GFP or STAT3 SER727ALA-GFP recombinant proteins in Hut78 STAT3 KDs electroporated with pLV-CMV-GFPSpark plasmid.

**Supplementary Figure 8.**
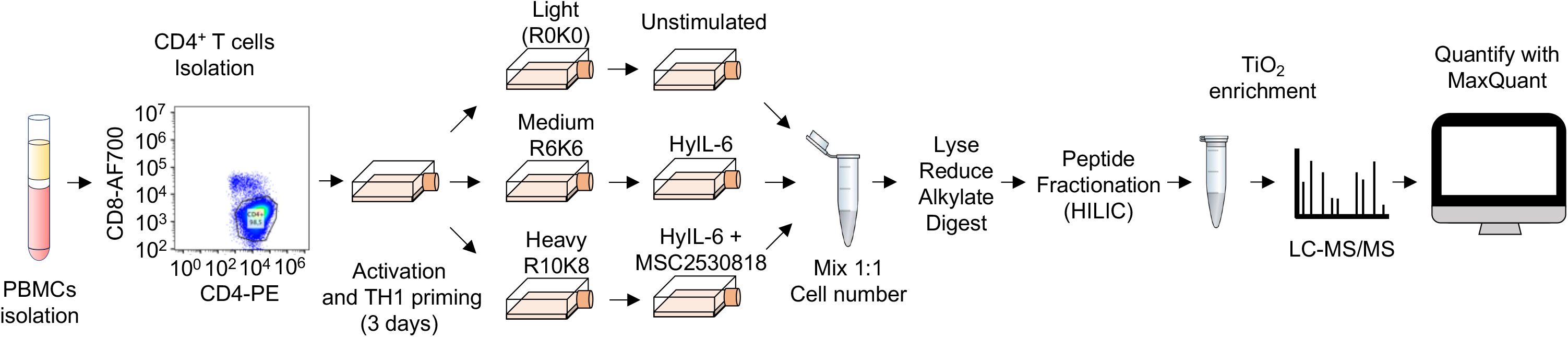
Role of CDK8 in the modulation of the HyIL-6-regulated phosphoproteome: Phosphoproteomics workflow. **A)** Experimental workflow of the phosphoproteomics experiment to elucidate the effect of CDK8 inhibition in the phosphoproteome regulated by HyIL-6 in human primary Th-1 cells.

**Supplementary Figure 9.**
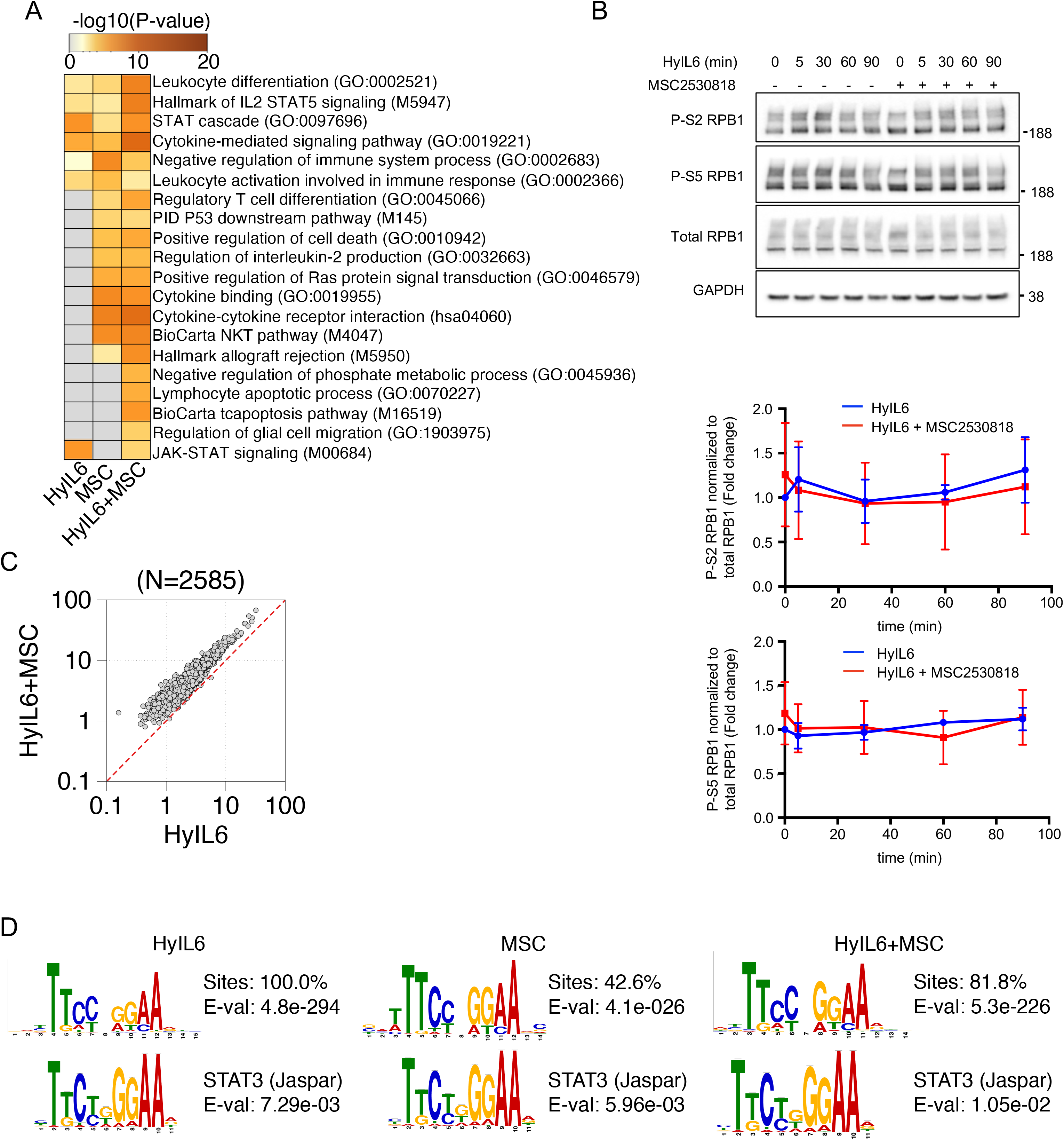
CDK8 fine-tunes STAT3 transcriptional program: **A)** Pathway analysis of differently expressed genes before and after HyIL6, MSC or HyIL6+MSC stimulation using Metascape (Zhou et al., 2019). Top 20 pathways are shown. B) Immunoblot analysis of the Ser2 and Ser5 phosphorylation state of RPB1 in Th-1 cells treated or untreated with MSC2530818 upon HyIL-6 stimulation. C) Shown are the most significant de novo motifs identified in STAT3 bound regions after HyIL6, MSC or HyIL6+MSC stimulation (top) and the matched STAT3 motif (bottom) from JASPAR database using TOMTOM.

## REFERENCES

Akamatsu, M., Mikami, N., Ohkura, N., Kawakami, R., Kitagawa, Y., Sugimoto, A., Hirota, K., Nakamura, N., Ujihara, S., Kurosaki, T., et al. (2019). Conversion of antigen-specific effector/memory T cells into Foxp3-expressing Treg cells by inhibition of CDK8/19. Sci Immunol 4.

Ali, A.K., Nandagopal, N., and Lee, S.H. (2015). IL-15-PI3K-AKT-mTOR: A Critical Pathway in the Life Journey of Natural Killer Cells. Front Immunol 6, 355.

Arai, K.I., Lee, F., Miyajima, A., Miyatake, S., Arai, N., and Yokota, T. (1990). Cytokines: coordinators of immune and inflammatory responses. Annu Rev Biochem 59, 783–836.

Bailey, T.L., Boden, M., Buske, F.A., Frith, M., Grant, C.E., Clementi, L., Ren, J., Li, W.W., and Noble, W.S. (2009). MEME SUITE: tools for motif discovery and searching. Nucleic Acids Res 37, W202–208.

Bancerek, J., Poss, Z.C., Steinparzer, I., Sedlyarov, V., Pfaffenwimmer, T., Mikulic, I., Dolken, L., Strobl, B., Muller, M., Taatjes, D.J., et al. (2013). CDK8 kinase phosphorylates transcription factor STAT1 to selectively regulate the interferon response. Immunity 38, 250–262.

Benjamini, Y., and Hochberg, Y. (1995). Controlling the False Discovery Rate - a Practical and Powerful Approach to Multiple Testing. J R Stat Soc B 57, 289–300.

Betz, U.A., and Muller, W. (1998). Regulated expression of gp130 and IL-6 receptor alpha chain in T cell maturation and activation. Int Immunol 10, 1175–1184.

Chen, M., Li, J., Liang, J., Thompson, Z.S., Kathrein, K., Broude, E.V., and Roninson, I.B. (2019). Systemic Toxicity Reported for CDK8/19 Inhibitors CCT251921 and MSC2530818 Is Not Due to Target Inhibition. Cells 8.

Chen, M., Liang, J., Ji, H., Yang, Z., Altilia, S., Hu, B., Schronce, A., McDermott, M.S.J., Schools, G.P., Lim, C.U., et al. (2017). CDK8/19 Mediator kinases potentiate induction of transcription by NFkappaB. Proc Natl Acad Sci U S A 114, 10208–10213.

Chung, J., Uchida, E., Grammer, T.C., and Blenis, J. (1997). STAT3 serine phosphorylation by ERK-dependent and -independent pathways negatively modulates its tyrosine phosphorylation. Mol Cell Biol 17, 6508–6516.

Cox, J., and Mann, M. (2008). MaxQuant enables high peptide identification rates, individualized p.p.b.-range mass accuracies and proteome-wide protein quantification. Nat Biotechnol 26, 1367–1372.

Cox, J., Neuhauser, N., Michalski, A., Scheltema, R.A., Olsen, J.V., and Mann, M. (2011). Andromeda: a peptide search engine integrated into the MaxQuant environment. J Proteome Res 10, 1794–1805.

Czodrowski, P., Mallinger, A., Wienke, D., Esdar, C., Poschke, O., Busch, M., Rohdich, F., Eccles, S.A., Ortiz-Ruiz, M.J., Schneider, R., et al. (2016). Structure-Based Optimization of Potent, Selective, and Orally Bioavailable CDK8 Inhibitors Discovered by High-Throughput Screening. J Med Chem 59, 9337–9349.

Debidda, M., Wang, L., Zang, H., Poli, V., and Zheng, Y. (2005). A role of STAT3 in Rho GTPase-regulated cell migration and proliferation. J Biol Chem 280, 17275–17285.

Decker, T., and Kovarik, P. (2000). Serine phosphorylation of STATs. Oncogene 19, 2628–2637.

Donner, A.J., Ebmeier, C.C., Taatjes, D.J., and Espinosa, J.M. (2010). CDK8 is a positive regulator of transcriptional elongation within the serum response network. Nat Struct Mol Biol 17, 194–201.

Donner, A.J., Szostek, S., Hoover, J.M., and Espinosa, J.M. (2007). CDK8 is a stimulus-specific positive coregulator of p53 target genes. Mol Cell 27, 121–133.

Fielding, C.A., McLoughlin, R.M., McLeod, L., Colmont, C.S., Najdovska, M., Grail, D., Ernst, M., Jones, S.A., Topley, N., and Jenkins, B.J. (2008). IL-6 regulates neutrophil trafficking during acute inflammation via STAT3. J Immunol 181, 2189–2195.

Fischer, M., Goldschmitt, J., Peschel, C., Brakenhoff, J.P., Kallen, K.J., Wollmer, A., Grotzinger, J., and Rose-John, S. (1997). I. A bioactive designer cytokine for human hematopoietic progenitor cell expansion. Nat Biotechnol 15, 142–145.

Fredriksson, S., Gullberg, M., Jarvius, J., Olsson, C., Pietras, K., Gustafsdottir, S.M., Ostman, A., and Landegren, U. (2002). Protein detection using proximity-dependent DNA ligation assays. Nat Biotechnol 20, 473–477.

Gabay, C. (2006). Interleukin-6 and chronic inflammation. Arthritis Res Ther 8 Suppl 2, S3.

Gorby, C., Martinez-Fabregas, J., Wilmes, S., and Moraga, I. (2018). Mapping Determinants of Cytokine Signaling via Protein Engineering. Front Immunol 9, 2143.

Guo, Z., Wang, G., Lv, Y., Wan, Y.Y., and Zheng, J. (2019). Inhibition of Cdk8/Cdk19 Activity Promotes Treg Cell Differentiation and Suppresses Autoimmune Diseases. Front Immunol 10, 1988.

Gupta, S., Stamatoyannopoulos, J.A., Bailey, T.L., and Noble, W.S. (2007). Quantifying similarity between motifs. Genome Biol 8, R24.

Hakemi, M.G., Ghaedi, K., Andalib, A., Hosseini, M., and Rezaei, A. (2011). Optimization of human Th17 cell differentiation in vitro: evaluating different polarizing factors. In Vitro Cell Dev-An 47, 581–592.

Han, J.M., Patterson, S.J., and Levings, M.K. (2012). The Role of the PI3K Signaling Pathway in CD4(+) T Cell Differentiation and Function. Front Immunol 3, 245.

Harlen, K.M., and Churchman, L.S. (2017). The code and beyond: transcription regulation by the RNA polymerase II carboxy-terminal domain. Nat Rev Mol Cell Biol 18, 263–273.

Heinrich, P.C., Behrmann, I., Muller-Newen, G., Schaper, F., and Graeve, L. (1998). Interleukin-6-type cytokine signalling through the gp130/Jak/STAT pathway. Biochem J 334 (Pt 2), 297–314.

Howden, A.J.M., Hukelmann, J.L., Brenes, A., Spinelli, L., Sinclair, L.V., Lamond, A.I., and Cantrell, D.A. (2019). Quantitative analysis of T cell proteomes and environmental sensors during T cell differentiation. Nat Immunol 20, 1542–1554.

Hu, W., Wang, G., Huang, D., Sui, M., and Xu, Y. (2019). Cancer Immunotherapy Based on Natural Killer Cells: Current Progress and New Opportunities. Front Immunol 10, 1205.

Huang da, W., Sherman, B.T., and Lempicki, R.A. (2009). Systematic and integrative analysis of large gene lists using DAVID bioinformatics resources. Nat Protoc 4, 44–57.

Huang, D.W., Sherman, B.T., Tan, Q., Collins, J.R., Alvord, W.G., Roayaei, J., Stephens, R., Baseler, M.W., Lane, H.C., and Lempicki, R.A. (2007). The DAVID Gene Functional Classification Tool: a novel biological module-centric algorithm to functionally analyze large gene lists. Genome Biol 8, R183.

Hunter, C.A., and Jones, S.A. (2015). IL-6 as a keystone cytokine in health and disease. Nat Immunol 16, 448–457.

Hutchins, A.P., Diez, D., and Miranda-Saavedra, D. (2013). Genomic and computational approaches to dissect the mechanisms of STAT3’s universal and cell type-specific functions. JAKSTAT 2, e25097.

Jeronimo, C., and Robert, F. (2017). The Mediator Complex: At the Nexus of RNA Polymerase II Transcription. Trends Cell Biol 27, 765–783.

Jonchere, B., Belanger, A., Guette, C., Barre, B., and Coqueret, O. (2013). STAT3 as a new autophagy regulator. JAKSTAT 2, e24353.

Jones, G.W., McLoughlin, R.M., Hammond, V.J., Parker, C.R., Williams, J.D., Malhotra, R., Scheller, J., Williams, A.S., Rose-John, S., Topley, N., et al. (2010). Loss of CD4+ T cell IL-6R expression during inflammation underlines a role for IL-6 trans signaling in the local maintenance of Th17 cells. J Immunol 184, 2130–2139.

Jones, S.A., and Jenkins, B.J. (2018). Recent insights into targeting the IL-6 cytokine family in inflammatory diseases and cancer. Nat Rev Immunol 18, 773–789.

Kim, H., and Baumann, H. (1997). The carboxyl-terminal region of STAT3 controls gene induction by the mouse haptoglobin promoter. J Biol Chem 272, 14571–14579.

Knuesel, M.T., Meyer, K.D., Bernecky, C., and Taatjes, D.J. (2009). The human CDK8 subcomplex is a molecular switch that controls Mediator coactivator function. Genes Dev 23, 439–451.

Korn, T., Mitsdoerffer, M., Croxford, A.L., Awasthi, A., Dardalhon, V.A., Galileos, G., Vollmar, P., Stritesky, G.L., Kaplan, M.H., Waisman, A., et al. (2008). IL-6 controls Th17 immunity in vivo by inhibiting the conversion of conventional T cells into Foxp3+ regulatory T cells. Proc Natl Acad Sci U S A 105, 18460–18465.

Kosciuczuk, E.M., Mehrotra, S., Saleiro, D., Kroczynska, B., Majchrzak-Kita, B., Lisowski, P., Driehaus, C., Rogalska, A., Turner, A., Lienhoop, T., et al. (2019). Sirtuin 2-mediated deacetylation of cyclin-dependent kinase 9 promotes STAT1 signaling in type I interferon responses. J Biol Chem 294, 827–837.

Krutzik, P.O., and Nolan, G.P. (2006). Fluorescent cell barcoding in flow cytometry allows high-throughput drug screening and signaling profiling. Nat Methods 3, 361–368.

Langmead, B., Trapnell, C., Pop, M., and Salzberg, S.L. (2009). Ultrafast and memory-efficient alignment of short DNA sequences to the human genome. Genome Biol 10, R25.

Laplante, M., and Sabatini, D.M. (2012). mTOR signaling in growth control and disease. Cell 149, 274–293.

LaPorte, S.L., Juo, Z.S., Vaclavikova, J., Colf, L.A., Qi, X., Heller, N.M., Keegan, A.D., and Garcia, K.C. (2008). Molecular and structural basis of cytokine receptor pleiotropy in the interleukin-4/13 system. Cell 132, 259–272.

Leonard, W.J., and Lin, J.X. (2000). Cytokine receptor signaling pathways. J Allergy Clin Immunol 105, 877–888.

Levy, D.E., and Darnell, J.E., Jr. (2002). Stats: transcriptional control and biological impact. Nat Rev Mol Cell Biol 3, 651–662.

Li, B., and Dewey, C.N. (2011). RSEM: accurate transcript quantification from RNA-Seq data with or without a reference genome. BMC Bioinformatics 12, 323.

Li, H., Handsaker, B., Wysoker, A., Fennell, T., Ruan, J., Homer, N., Marth, G., Abecasis, G., Durbin, R., and Genome Project Data Processing, S. (2009). The Sequence Alignment/Map format and SAMtools. Bioinformatics 25, 2078–2079.

Lim, C.P., and Cao, X. (1999). Serine phosphorylation and negative regulation of Stat3 by JNK. J Biol Chem 274, 31055–31061.

Lin, J.X., and Leonard, W.J. (2019). Fine-Tuning Cytokine Signals. Annu Rev Immunol 37, 295–324.

Liu, B., Palmfeldt, J., Lin, L., Colaco, A., Clemmensen, K.K.B., Huang, J., Xu, F., Liu, X., Maeda, K., Luo, Y., et al. (2018). STAT3 associates with vacuolar H(+)-ATPase and regulates cytosolic and lysosomal pH. Cell Res 28, 996–1012.

Liu, Q., Kirubakaran, S., Hur, W., Niepel, M., Westover, K., Thoreen, C.C., Wang, J., Ni, J., Patricelli, M.P., Vogel, K., et al. (2012). Kinome-wide selectivity profiling of ATP-competitive mammalian target of rapamycin (mTOR) inhibitors and characterization of their binding kinetics. J Biol Chem 287, 9742–9752.

Liu, S.T., Pham, H., Pandol, S.J., and Ptasznik, A. (2013). Src as the link between inflammation and cancer. Front Physiol 4, 416.

Lloyd-Lewis, B., Krueger, C.C., Sargeant, T.J., D’Angelo, M.E., Deery, M.J., Feret, R., Howard, J.A., Lilley, K.S., and Watson, C.J. (2018). Stat3-mediated alterations in lysosomal membrane protein composition. J Biol Chem 293, 4244–4261.

Luke, J.J., D’Adamo, D.R., Dickson, M.A., Keohan, M.L., Carvajal, R.D., Maki, R.G., de Stanchina, E., Musi, E., Singer, S., and Schwartz, G.K. (2012). The cyclin-dependent kinase inhibitor flavopiridol potentiates doxorubicin efficacy in advanced sarcomas: preclinical investigations and results of a phase I dose-escalation clinical trial. Clin Cancer Res 18, 2638–2647.

Martinez-Fabregas, J., Prescott, A., van Kasteren, S., Pedrioli, D.L., McLean, I., Moles, A., Reinheckel, T., Poli, V., and Watts, C. (2018). Lysosomal protease deficiency or substrate overload induces an oxidative-stress mediated STAT3-dependent pathway of lysosomal homeostasis. Nature Communications 9.

Martinez-Fabregas, J., Wilmes, S., Wang, L.P., Hafer, M., Pohler, E., Lokau, J., Garbers, C., Cozzani, A., Fyfe, P.K., Piehler, J., et al. (2019). Kinetics of cytokine receptor trafficking determine signaling and functional selectivity. Elife 8.

Mauer, J., Denson, J.L., and Bruning, J.C. (2015). Versatile functions for IL-6 in metabolism and cancer. Trends Immunol 36, 92–101.

Miyahara, Y., Odunsi, K., Chen, W.H., Peng, G.Y., Matsuzaki, J., and Wang, R.F. (2008). Generation and regulation of human CD4+ IL-17-producing T cells in ovarian cancer. P Natl Acad Sci USA 105, 15505–15510.

Ng, D.C., Lin, B.H., Lim, C.P., Huang, G., Zhang, T., Poli, V., and Cao, X. (2006). Stat3 regulates microtubules by antagonizing the depolymerization activity of stathmin. J Cell Biol 172, 245–257.

O’Shea, J.J., and Murray, P.J. (2008). Cytokine signaling modules in inflammatory responses. Immunity 28, 477–487.

Pilz, A., Ramsauer, K., Heidari, H., Leitges, M., Kovarik, P., and Decker, T. (2003). Phosphorylation of the Stat1 transactivating domain is required for the response to type I interferons. EMBO Rep 4, 368–373.

Poli, V., and Camporeale, A. (2015). STAT3-Mediated Metabolic Reprograming in Cellular Transformation and Implications for Drug Resistance. Front Oncol 5, 121.

Putz, E.M., Gotthardt, D., Hoermann, G., Csiszar, A., Wirth, S., Berger, A., Straka, E., Rigler, D., Wallner, B., Jamieson, A.M., et al. (2013). CDK8-mediated STAT1-S727 phosphorylation restrains NK cell cytotoxicity and tumor surveillance. Cell Rep 4, 437–444.

Quinlan, A.R., and Hall, I.M. (2010). BEDTools: a flexible suite of utilities for comparing genomic features. Bioinformatics 26, 841–842.

Ramirez, F., Ryan, D.P., Gruning, B., Bhardwaj, V., Kilpert, F., Richter, A.S., Heyne, S., Dundar, F., and Manke, T. (2016). deepTools2: a next generation web server for deep-sequencing data analysis. Nucleic Acids Res 44, W160–165.

Rhee, S.H., Jones, B.W., Toshchakov, V., Vogel, S.N., and Fenton, M.J. (2003). Toll-like receptors 2 and 4 activate STAT1 serine phosphorylation by distinct mechanisms in macrophages. J Biol Chem 278, 22506–22512.

Ridgley, L.A., Anderson, A.E., Maney, N.J., Naamane, N., Skelton, A.J., Lawson, C.A., Emery, P., Isaacs, J.D., Carmody, R.J., and Pratt, A.G. (2019). IL-6 Mediated Transcriptional Programming of Naive CD4+ T Cells in Early Rheumatoid Arthritis Drives Dysregulated Effector Function. Front Immunol 10, 1535.

Robinson, J.T., Thorvaldsdottir, H., Winckler, W., Guttman, M., Lander, E.S., Getz, G., and Mesirov, J.P. (2011). Integrative genomics viewer. Nat Biotechnol 29, 24–26.

Robinson, M.D., McCarthy, D.J., and Smyth, G.K. (2010). edgeR: a Bioconductor package for differential expression analysis of digital gene expression data. Bioinformatics 26, 139–140.

Rose-John, S. (2012). IL-6 trans-signaling via the soluble IL-6 receptor: importance for the pro-inflammatory activities of IL-6. Int J Biol Sci 8, 1237–1247.

Rose-John, S. (2018). Interleukin-6 Family Cytokines. Cold Spring Harb Perspect Biol 10.

Ross, S.H., Rollings, C., Anderson, K.E., Hawkins, P.T., Stephens, L.R., and Cantrell, D.A. (2016). Phosphoproteomic Analyses of Interleukin 2 Signaling Reveal Integrated JAK Kinase-Dependent and -Independent Networks in CD8(+) T Cells. Immunity 45, 685–700.

Sadzak, I., Schiff, M., Gattermeier, I., Glinitzer, R., Sauer, I., Saalmuller, A., Yang, E., Schaljo, B., and Kovarik, P. (2008). Recruitment of Stat1 to chromatin is required for interferon-induced serine phosphorylation of Stat1 transactivation domain. Proc Natl Acad Sci U S A 105, 8944–8949.

Scheller, J., Chalaris, A., Schmidt-Arras, D., and Rose-John, S. (2011). The pro- and antiinflammatory properties of the cytokine interleukin-6. Biochim Biophys Acta 1813, 878–888.

Schindler, C., Levy, D.E., and Decker, T. (2007). JAK-STAT signaling: from interferons to cytokines. J Biol Chem 282, 20059–20063.

Sekiya, T., and Yoshimura, A. (2016). In Vitro Th Differentiation Protocol. Methods Mol Biol 1344, 183–191.

Servais, F.A., Kirchmeyer, M., Hamdorf, M., Minoungou, N.W.E., Rose-John, S., Kreis, S., Haan, C., and Behrmann, I. (2019). Modulation of the IL-6-Signaling Pathway in Liver Cells by miRNAs Targeting gp130, JAK1, and/or STAT3. Mol Ther Nucleic Acids 16, 419–433.

Shah, M., Patel, K., Mukhopadhyay, S., Xu, F., Guo, G., and Sehgal, P.B. (2006). Membrane-associated STAT3 and PY-STAT3 in the cytoplasm. J Biol Chem 281, 7302–7308.

Silva, C.M. (2004). Role of STATs as downstream signal transducers in Src family kinase-mediated tumorigenesis. Oncogene 23, 8017–8023.

Smith-Garvin, J.E., Koretzky, G.A., and Jordan, M.S. (2009). T cell activation. Annu Rev Immunol 27, 591–619.

Steen, H.C., Kotredes, K.P., Nogusa, S., Harris, M.Y., Balachandran, S., and Gamero, A.M. (2016). Phosphorylation of STAT2 on serine-734 negatively regulates the IFN-alpha-induced antiviral response. J Cell Sci 129, 4190–4199.

Stroud, R.M., and Wells, J.A. (2004). Mechanistic diversity of cytokine receptor signaling across cell membranes. Sci STKE 2004, re7.

Subramanian, A., Tamayo, P., Mootha, V.K., Mukherjee, S., Ebert, B.L., Gillette, M.A., Paulovich, A., Pomeroy, S.L., Golub, T.R., Lander, E.S., et al. (2005). Gene set enrichment analysis: a knowledge-based approach for interpreting genome-wide expression profiles. Proc Natl Acad Sci U S A 102, 15545–15550.

Takeda, K., Noguchi, K., Shi, W., Tanaka, T., Matsumoto, M., Yoshida, N., Kishimoto, T., and Akira, S. (1997). Targeted disruption of the mouse Stat3 gene leads to early embryonic lethality. Proc Natl Acad Sci U S A 94, 3801–3804.

Tanaka, T., Narazaki, M., and Kishimoto, T. (2014). IL-6 in inflammation, immunity, and disease. Cold Spring Harb Perspect Biol 6, a016295.

Teng, T.S., Lin, B., Manser, E., Ng, D.C., and Cao, X. (2009). Stat3 promotes directional cell migration by regulating Rac1 activity via its activator betaPIX. J Cell Sci 122, 4150–4159.

Thoreen, C.C., Kang, S.A., Chang, J.W., Liu, Q., Zhang, J., Gao, Y., Reichling, L.J., Sim, T., Sabatini, D.M., and Gray, N.S. (2009). An ATP-competitive mammalian target of rapamycin inhibitor reveals rapamycin-resistant functions of mTORC1. J Biol Chem 284, 8023–8032.

Varinou, L., Ramsauer, K., Karaghiosoff, M., Kolbe, T., Pfeffer, K., Muller, M., and Decker, T. (2003). Phosphorylation of the Stat1 transactivation domain is required for full-fledged IFN-gamma-dependent innate immunity. Immunity 19, 793–802.

Wang, X., Lupardus, P., Laporte, S.L., and Garcia, K.C. (2009). Structural biology of shared cytokine receptors. Annu Rev Immunol 27, 29–60.

Wen, Z., Zhong, Z., and Darnell, J.E., Jr. (1995). Maximal activation of transcription by Stat1 and Stat3 requires both tyrosine and serine phosphorylation. Cell 82, 241–250.

Wilmes, S., Hafer, M., Vuorio, J., Tucker, J.A., Winkelmann, H., Lochte, S., Stanly, T.A., Pulgar Prieto, K.D., Poojari, C., Sharma, V., et al. (2020). Mechanism of homodimeric cytokine receptor activation and dysregulation by oncogenic mutations. Science 367, 643–652.

Witalisz-Siepracka, A., Gotthardt, D., Prchal-Murphy, M., Didara, Z., Menzl, I., Prinz, D., Edlinger, L., Putz, E.M., and Sexl, V. (2018). nK Cell-Specific CDK8 Deletion Enhances Antitumor Responses. Cancer Immunol Res 6, 458–466.

Yang, J., Kunimoto, H., Katayama, B., Zhao, H., Shiromizu, T., Wang, L., Ozawa, T., Tomonaga, T., Tsuruta, D., and Nakajima, K. (2020). Phospho-Ser727 triggers a multistep inactivation of STAT3 by rapid dissociation of pY705-SH2 through C-terminal tail modulation. Int Immunol 32, 73–88.

Yokogami, K., Wakisaka, S., Avruch, J., and Reeves, S.A. (2000). Serine phosphorylation and maximal activation of STAT3 during CNTF signaling is mediated by the rapamycin target mTOR. Curr Biol 10, 47–50.

Zhang, W., and Liu, H.T. (2002). MAPK signal pathways in the regulation of cell proliferation in mammalian cells. Cell Res 12, 9–18.

Zhang, Y., Liu, T., Meyer, C.A., Eeckhoute, J., Johnson, D.S., Bernstein, B.E., Nusbaum, C., Myers, R.M., Brown, M., Li, W., et al. (2008). Model-based analysis of ChIP-Seq (MACS). Genome Biol 9, R137.

Zhou, Y., Zhou, B., Pache, L., Chang, M., Khodabakhshi, A.H., Tanaseichuk, O., Benner, C., and Chanda, S.K. (2019). Metascape provides a biologist-oriented resource for the analysis of systems-level datasets. Nat Commun 10, 1523.

